# Caveolin-1–Mediated Blood-brain Barrier Transcytosis Promotes *Porphyromonas gingivalis* Invasion and Alzheimer’s Disease-Like Changes

**DOI:** 10.64898/2026.02.17.706258

**Authors:** Chenghan Mei, Qixing Zhong, Xiaojie Li, Yao He, Ting Zhang, Bangcheng Zhao, Jiangshang Liu, Zhihui Zhong

## Abstract

Pathogens such as *Porphyromonas gingivalis* (*P. gingivalis*) may lead to Alzheimer’s disease (AD), but how they get into the brain and cause AD remains unclear, especially their level in blood is usually low. AD involves early breakdown of the blood-brain barrier (BBB). BBB disruption may allow blood-derived neurotoxic components to penetrate the BBB and enter the brain. We investigated whether BBB disruption permits *P. gingivalis* to enter the brain and accelerate disease-related changes. Using animal models, human BBB organoids and bEnd.3 cells, we found that a leaky BBB, especially through increased caveolin-1-dependent transcytosis, allowed *P. gingivalis* to cross into the brain. The bacteria triggered the hyperphosphorylation of Tau protein, increased Aβ levels, and sustained neuroinflammation via activated glial cells. These findings indicate that BBB dysfunction facilitates *P. gingivalis* invasion, creating a vicious cycle of infection and neurodegeneration that may drive AD progression.

## Introduction

Alzheimer’s disease (AD)—a progressive neurodegenerative disorder characterized by amyloid plaques, neurofibrillary tangles (NFTs), and neuroinflammation—imposes substantial socioeconomic burdens on families and healthcare systems amid escalating global prevalence. The amyloid cascade hypothesis remains the prevailing etiological framework. To date, only two monoclonal antibodies targeting amyloid-β protein (Aβ), lecanemab and donanemab, have been approved for AD treatment [1]. A growing consensus indicates that impaired Aβ clearance and overproduction constitute an early initiating event in AD pathogenesis [2]. Mounting evidence supports the pathogen hypothesis of AD pathogenesis. Microbial infections—including Herpes simplex virus-1 (HSV-1), *Chlamydia pneumoniae*, *Borrelia burgdorferi*, and *Porphyromonas gingivalis* (*P. gingivalis*)—trigger elevated cerebral Aβ production and deposition, implicating innate immune responses in disease initiation [3, 4]. Aβ is a type of antimicrobial peptide that can fight pathogens by aggregating and encapsulating them. However, insufficient clearance of Aβ intensifies neuroinflammation, thereby perpetuating a vicious cycle of Aβ deposition–inflammatory response [5].

Oral dysbiosis is closely associated with the development of AD, which may lead to oral bacteria entering the bloodstream and spreading to the brain. Oral bacterial infections may therefore be one of the potential causes of AD [6]. Noble et al. (2014) reported that elevated levels of serum IgG against common periodontal pathogens were associated with an increased risk of developing AD [7]. *P. gingivalis* is one of the key pathogens involved in the development of periodontitis and is associated with the deposition of Aβ and the pathogenesis of AD. Notably, *P. gingivalis* DNA has been detected in the cerebrospinal fluid (CSF) and saliva of AD patients, and levels of *P. gingivalis*-specific proteases, known as gingipains, in AD patient brain tissue correlate with Tau protein pathology [4]. Additionally, infection with *P. gingivalis* resulted in a significant increase in hippocampal Aβ deposition in APP-Tg mice [8]. In the middle-aged mice infected with *P. gingivalis*, the levels of TNF-α, IL-6, and IL-1β were upregulated in the brain [9]. When mice were infected with *P. gingivalis*, the influx of Aβ increased, and it was found to localise around cerebral endothelial cells [10]. The deposition of Aβ around cerebral blood vessels can trigger cerebral amyloid angiopathy (CAA) [11]. Moreover, CAA induces cerebrovascular dysfunction and degeneration, directly driving cognitive impairment and potentially representing a pathological nexus between vascular dementia and AD [12].

Although many studies have reported a correlation between *P. gingivalis* and AD, and have shown that it can induce memory impairment and exacerbate AD features [8, 9], the mechanism by which *P. gingivalis* invades the brain remains incompletely elucidated. *P. gingivalis* may enter the brain via the nerves, lymphatic system, or blood system. Although the brain has no lymphatic system, it does have a glymphatic system that transfers CSF and brain interstitial fluid, facilitating the elimination of metabolic waste products from the brain [13]. *P. gingivalis* may invade the brain through a compromised blood-brain barrier (BBB). Lei Shuang et al. (2023) demonstrated that prolonged systemic administration of *P. gingivalis* at supraphysiological doses (1 × 10⁸ colony-forming unit, CFU) induces BBB disruption in rat hippocampal and cortical regions, enabling brain infiltration by the pathogen [14]. While these models demonstrate BBB disruption by *P. gingivalis*, the non-physiological dosing undermines the relevance of these findings to the bacterium’s ability to infiltrate the BBB physiologically.

The BBB is crucial for brain function and consists of tight and adherens junctions. Under normal conditions, it prevents blood cells and pathogens from infiltrating the brain [15]. The BBB also regulates the entry and exit of molecules into and out of the brain, thereby tightly controlling the brain’s chemical environment. BBB permeability increased before cognitive decline in AD [16]. Mild cognitive impairment (MCI) is an intermediate state between normal ageing and AD. BBB permeability in the hippocampus increased significantly in patients with MCI [17]. As the hippocampus is the first brain region to be affected by AD, it exhibits synaptic loss and increased BBB permeability during early stages of MCI and AD. Hippocampal BBB permeability increases with age, with aging itself being a risk factor for AD, and this increase is further heightened in individuals with MCI [18]. The apolipoprotein E4 (*APOE4*) gene is the strongest genetic risk factor for sporadic AD. *APOE4* promotes the accelerated breakdown of the BBB and the degeneration of brain capillary pericytes [19]. Most importantly, deposits of aberrant bacterial lipopolysaccharide (LPS) have been observed around the blood vessels of the brain in patients with AD [20], suggesting that the breakdown of the BBB may facilitate the entry of toxic blood-derived molecules, cells, and microbial pathogens into the brain, thereby triggering neuroinflammatory and immune responses.

In previous studies, we found that the BBB is susceptible to damage in various animal models of vascular lesions. Longitudinal studies in rhesus monkeys with middle cerebral artery occlusion (MCAO) documented persistent BBB compromise, with significantly elevated bilateral hippocampal BBB permeability persisting >12 months post-infarct [21]. Additionally, the long-term MCAO monkeys demonstrated abnormal Aβ metabolic profiles [21], suggesting that cerebrovascular injury could contribute to AD pathology by disrupting Aβ metabolism. Chronic cerebral ischemia is one of the causes of vascular dementia (VD) and AD [22, 23]. VD and AD share complex and overlapping pathophysiological associations. In most dementia patients, particularly the elderly, mixed pathology—coexisting AD and VD pathologies—is frequently observed [24]. Moreover, bilateral common carotid artery occlusion (2VO) is a well-established rat model for studying chronic cerebral ischemia. Jae-Hyung Park et al (2019) reported that chronic cerebral hypoperfusion in 2VO rats resulted in elevated Aβ and p-tau expression alongside selectively reduced hippocampal neuronal activity [25]. Impairment of BBB integrity is a feature shared by both AD and VD. Both our previous studies and those reported by other groups have shown that 2VO rats develop increased BBB permeability after surgery [26, 27]. Furthermore, changes to the gut microbiota caused by antibiotic therapy can impair the integrity of the BBB in rhesus monkeys and mice [28, 29].

Based on previous studies, vascular lesions can induce BBB dysfunction and abnormal Aβ metabolism. Furthermore, cerebrovascular lesions are key risk factors for AD, with cerebrovascular dysfunction contributing significantly to the disease pathogenesis [30]. Therefore, it was of our interest to investigate whether vascular lesions leading to BBB dysfunction might permit specific pathogens to cross the barrier, thereby triggering an innate immune response that contributes to AD-like neurodegenerative changes. We hypothesize that individuals with compromised brain vasculature exhibit heightened susceptibility to neurotoxicity mediated by pathogens from the periphery. In this context, BBB leakage—particularly in the hippocampus—allows microbes to enter the brain, triggering innate immunity–driven Aβ pathology characteristic of AD. This study aims to investigate whether vascular injury–induced BBB disruption leads to specific pathogens passing through the BBB, thus triggering an innate immune response.

## Materials and Methods

### Experimental animals

4 adult male rhesus monkeys, aged 7-12 years and weighing 7-10 kg, were obtained from Sichuan Green-House Biotech Co., Ltd. The monkeys were individually housed in cages under a 12-hour light/dark cycle, at a temperature of 23 ± 2 °C, and with a humidity level of 50–70%. The monkeys were fed twice daily and had unrestricted access to water. No monkeys were sacrificed in this study. Male Wistar rats (n > 50), aged 4–6 weeks and weighing 320–340 g, were purchased from Beijing Charles River Laboratory Animal Technology Company (cat. 102). The rats were raised in a specific pathogen-free animal facility under a 12-hour light/dark cycle, at a temperature of 22 ± 1 °C and humidity of 50–60%. All experimental protocols were adhered to the ’Guidance Suggestions for the Care and Use of Laboratory Animals’ issued by the Ministry of Science and Technology of the People’s Republic of China, as well as the animal ethical criteria established by the Association for Assessment and Accreditation of Laboratory Animal Care. The protocols were approved by the Animal Care Committee of Sichuan University West China Hospital (approval number 2020119A).

### 2VO model establishment

In brief, rats were fasted for 3-4 hours before the operation, while free access to water was maintained. The rats were then anaesthetized with isoflurane, after which a 1.5–2 cm long midline incision was made along the ventral cervical region of each rat. The common carotid artery, nerves, and muscles were carefully dissected using forceps, after which the common carotid artery was ligated using a 5-0 silk ligature. In the SHAM group, the rats underwent the same procedure, except without ligation of the common carotid artery. The neck incision was then closed using a 5-0 silk suture, and the anaesthesia was discontinued. Body temperature was maintained at 37 °C with a heating pad throughout the surgery until recovery.

### Brain LPS measurement

The dose conversion calculations were conducted based on quantitative measurements of bacterial exposure concentrations in these clinical cases [31]. Rats were injected with 200 μL of saline containing 5,000 CFU of *P. gingivalis* via the tail vein for 3 days following surgery. After 72 hours, the rats were anaesthetised and transcardially perfused with 200 mL of pharmaceutical-grade saline. Their hippocampus and cortex were collected and weighed. The brain tissue was homogenised with LAL H₂O (85 mg/750 μL) using a Dounce homogeniser. The level of LPS in the brain tissue was measured using a ToxinSensor™ Chromogenic LAL Endotoxin Assay Kit (Genscript, #L00350, USA). After completing the procedure in accordance with the instructions, the absorbance was detected at 545 nm, after which the LPS level in the samples was calculated.

### Intracerebral bacterial tracing experiment in rats

Following the 2VO surgery, rats received daily tail vein injections of 5,000 CFU (500 CFU/mL) of *P. gingivalis* for 3 consecutive days. 6 hours after the final bacterial injection, CSF was extracted from the rats. 20 μL of CSF was spread evenly on a blood plate and cultured anaerobically at 37 °C. 10 days later, the blood plate was photographed, and representative bacterial colonies were extracted to isolate DNA using a magnetic universal genomic DNA kit (Tiangen, #DP705, China). PCR was performed using 2×Taq Plus Master Mix II (Vazyme, #P213-01, China) with *P. gingivalis* 16S rRNA primers (The sequences of the primers were listed in Table S1) to determine whether the bacterial colonies were *P. gingivalis*. The PCR products were then subjected to Sanger sequencing.

After surgery, rats were administered, through tail vein injection, of 5,000 CFU of Cy5-labeled *P. gingivalis* once daily for 3 consecutive days. On day 4, rats were anaesthetised, followed by CSF collection and transcardial perfusion with PBS. 10 μL of CSF were mixed with 0.5% agarose on microscope slides for imaging using a confocal fluorescence microscope (Olympus, SpinSR, Japan). The rats’ brains were then flash-frozen using isopentane and liquid nitrogen. The brain tissue was embedded using an optimal cutting temperature compound, and 8 µm coronal sections were cut using a cryostat (Leica, LEICA CM 1950, German). The sections were then incubated with tomato lectin (8 μg/mL). The nuclei were then counterstained with DAPI, and images were captured using a confocal fluorescence microscope (Olympus, SpinSR, Japan).

### Cerebral cortex blood flow measurement

After anesthesia with isoflurane and shaving of the head fur, a midline sagittal incision was made to expose the skull. The skull was then carefully thinned to translucency using a drill and cleaned with a cotton applicator and saline. The Laser Scattered Blood Flow Imaging System (Moor Instruments Ltd, moorO2Flo-2, UK) was used to monitor cerebral blood flow (CBF) before and after 2VO surgery (on days 1 and 3). The imaging parameters and conditions were maintained consistently across all time points. After CBF acquisition, the wound was sutured, and anaesthesia was withdrawn.

### Evaluating BBB integrity in rats using Evans blue

Evans blue (Sigma, #E2129, USA) was used to evaluate BBB permeability in rats on days 1, 2, and 3 post-surgery. In brief, 2% Evans blue (3 mL/kg) was injected intravenously through the tail vein and allowed to circulate for two hours. Rats were anaesthetised, followed by transcardial perfusion with PBS. The hippocampus and cortex were quickly collected and weighed. Then, 50% trichloroacetic acid (3 μL per mg of brain tissue) and two 3 mm steel balls were added to the brain tissue, and the sample was homogenised for 1 minute at 70 Hz. Centrifugation was then performed at 10,000 g for 20 minutes, after which 100 μL of the supernatant was mixed with 300 μL of 95% ethanol. 5 mg of Evans blue was dissolved in 50% trichloroacetic acid and then diluted stepwise to 4 μg/mL. This solution was then diluted with 95% ethanol to create standard curve samples at concentrations of 1600, 800, 400, 200, 100, 50, 25, and 12.5 ng/mL, respectively. 100 μL of the samples was then added to a 96-well plate, and the fluorescence absorption intensity of the samples was evaluated using a microplate reader with an excitation wavelength of 620 nm and an emission wavelength of 680 nm.

### *P. gingivalis* culture and labelling with Cy5-ADA

*P. gingivalis* W83 was obtained from the Guangdong Microbial Culture Collection Centre and cultured anaerobically at 37 °C in brain-heart infusion (BHI) medium (BD, #GD-237400-100g, USA) containing 5 μg/mL of hematin chloride (Shanghai Yuanye, #S19203, China) and 10 μg/mL of vitamin K (Shanghai Yuanye, # B21297, China). To label *P. gingivalis* with Cy5-ADA (Chinese peptide, #979937, China), Cy5-ADA was added to the BHI medium (200 μM), and the *P. gingivalis* (OD_600 nm_ = 0.3–0.4) was cultured for 6 hours. *P. gingivalis* in the logarithmic phase was then used for the subsequent experiments.

### Microvessels isolation from the hippocampus and cortex of rat

As described previously [32], rat hippocampal and cortical microvessels were isolated on days 1 and 3 after 2VO surgery. After the rats were anaesthetised, the hippocampus and cortex were separated on ice, with the large vessels and meninges removed. The brain tissue was then homogenised in cold DMEM (Gibco, #C11995500BT, USA). Cerebral microvessels were isolated by centrifugation at 4 °C (10,000 g) for 15 minutes using 15% dextran. A 40 μm strainer was then used to retrieve the microvessels, which were subsequently washed with cold 0.5% BSA DMEM medium. The suspension was then centrifuged at 4 °C (3,000 g) for 10 minutes. The microvessels were collected for further analysis after the supernatant had been removed.

### Transmission electron microscopy (TEM) analysis

On day 1 post-operation, the rats were anesthetized, and tissue samples from the cortex and hippocampus were collected and immediately fixed in 2.5% glutaraldehyde. The samples were then post-fixed in 1% osmium tetroxide, dehydrated through a graded series of acetone, and infiltrated and embedded in Epon 812 resin. Semithin sections were stained with methylene blue for preliminary assessment. Ultrathin sections were cut using a diamond knife, followed by double-staining with uranyl acetate and lead citrate. Finally, the ultrastructure of the brain tissues was examined under a TEM (JOEL, JEM-1400 FLASH, Japan).

### RNA isolation and RT-qPCR analysis

Total RNA was isolated from samples using TRIzol reagent. Following the manufacturer’s guidelines, cDNA was synthesised using the cDNA Synthesis Kit (Vazyme, # R323-01, China). RT-qPCR was performed using a CFX Connect instrument (Bio-Rad, CFX Duet Real-Time PCR System, USA) and the SYBR qPCR Master Mix (Vazyme, #Q711, China). The detailed RT-qPCR conditions are provided in Table S2. Gene mRNA expression levels were evaluated using the 2^(−ΔΔCT) method, and all primers are presented in Table S1.

### BBB organoid establishment

Primary human pericytes and astrocytes were obtained from Shanghai Xuanke Biotechnology Co., Ltd., and were grown in pericyte medium (Sciencell, #1201, USA) and astrocyte growth medium (Meisen Cell, # CTCC-007-PriMed, China), respectively. hCMEC/D3 cells were purchased from Shanghai Jinyuan Biotechnology Co., Ltd. and cultured in EGM-2 Endothelial Basal Medium (Lonza, #CC-3162, USA). The cells were cultured in T25 or T75 flasks. The pericytes and astrocytes were maintained between passages 2 and 5, and the hCMEC/D3 cells between passages 2 and 10. The cells were detached using 0.25% trypsin-EDTA and resuspended in BBB working medium (EGM-2 medium without VEGF and FBS, containing 2% human serum). The cells were counted and then adjusted to an appropriate concentration of 1:1:1 ratio using the BBB working medium. The mixed cells were seeded in a low-adhesion agarose 96-well plate [33]. Each microwell contained 1500 cells of each type. The cells were then incubated at 37 °C with 5% CO₂ for 48 hours.

The oxygen-glucose deprivation (OGD) modelling was conducted as follows: 3 days after the BBB organoids were established, 300 μL of Earle’s balanced salt solution (Shanghai Yuanye, # R20055-1L, China) was added to the medium, and the plate was placed in a hypoxic chamber conditioned with 5% CO₂, 95% N₂ (10 L/minutes) until the oxygen concentration was below 1%. In the control group, the medium was replaced with the BBB working medium. The BBB organoids were cultured for 48 hours for subsequent analyses.

To evaluate the permeability of the BBB organoids, 100 μL of BBB working medium containing 5 μg/mL Cy5-albumin was added to the culture medium and maintained in darkness at 37 °C with 5% CO₂ for 12 hours. Then, 1 μL of 2 mg/mL Hoechst 33258 (Biosharp, # BL804A, China) was added to the medium, and the culture was incubated at 37°C for 30 minutes. The BBB organoids were then washed three times with saline and fixed with 4% paraformaldehyde. The BBB organoids were placed in an 8-well chamber slide for imaging using a confocal fluorescence microscope (Olympus, SpinSR, Japan). The mean fluorescence intensity of Cy5-albumin within the BBB organoids was measured at a depth of 50 µm using ImageJ (NIH, 1.54f, USA).

After OGD modelling, the BBB organoids were co-cultured with Cy5-ADA-labelled *P. gingivalis* at multiplicity of infection (MOI) 1 or 10 in the dark at 37 °C with 5% CO₂ for 12 hours. Then, 1 μL of 2 mg/mL Hoechst 33258 (Biosharp, # BL804A, China) was added to the medium, and the organoids were cultured for an additional 30 minutes. The BBB organoids were washed three times with saline and subsequently fixed using 4% paraformaldehyde. The organoids were then transferred to an 8-well chamber slide for imaging using a confocal fluorescence microscope (Olympus, SpinSR, Japan). The mean fluorescence intensity of the maximum intensity projection was generated from the bottom of the BBB organoid to a depth of 50 µm.

### Human brain organoids (BOs) establishment

The human BOs were established from H9 human embryonic stem cells (ESCs) (Beina Biotechnology Co., LTD., China) in accordance with the manufacturer’s guidelines (STEMdiff™, #80570, Canada). The detailed culture procedure was the same as in our previous study [34]. After culturing the BOs for more than 60 days, they were harvested for subsequent analyses. For coculturing with *P. gingivalis*, the BOs (73 days) were transferred to a 24-well ultra-low attachment plate, and 700 μL maturation medium was added per well. *P. gingivalis* were added to the well (0, 15,000, 150,000 CFU) and cultured at 37℃ with 5% CO_2_ for 72 hours. The BOs and culture medium were then collected for subsequent experiments.

### RNA sequencing (RNA-seq) and data analysis

The BOs (73 days) were harvested, and the RNAs were extracted. RNA purity and quantification were measured with a NanoDrop2000 spectrophotometer (Thermo Scientific, USA), and integrity was evaluated using an Agilent 2100 Bioanalyzer (Agilent Technologies, USA). Libraries were then constructed with the VAHTS Universal V10 RNA-seq Library Prep Kit (Premixed Version) following the manufacturer’s protocol. RNA sequencing was performed using the Illumina Novaseq 6000 platform with an average of 24 million reads per run. Cell-type composition was estimated using CIBERSORT [35]. Brain cell-type signatures (Cell Atlas) were obtained from the study by Gavin Sutton et al [36].

### bEnd.3 cell culture and OGD modelling

The bEnd.3 mouse cerebral microvascular endothelial cell line was obtained from the China National Collection of Authenticated Cell Cultures and maintained in DMEM medium (Gibco, #10566016, USA) containing 10% FBS. bEnd.3 cells were seeded in 6 or 12-well plates. For transfection, the cells were seeded in a 12-well plate. Mouse *Caveolin-1* (*Cav-1*) siRNA and NC siRNA were purchased from GenePharma, and siRNA Mete Plus transfection reagent (GenePharma, #G04026, China) was used for delivery. The detailed siRNA sequence was presented in Table S3. After overnight culturing, *Cav-1* or NC siRNA was transfected into the bEnd.3 cells for 30 hours.

OGD modelling was performed to mimic acute ischaemic conditions in bEnd.3 cells. Briefly, bEnd.3 cells were cultured as a monolayer in DMEM medium containing 1% FBS overnight. The cell medium was then changed to glucose-free DMEM without FBS. The cells were then placed in a hypoxic chamber at 37 °C with a constant N₂ flow of 5 L/minutes for 2.5 hours. Regarding the control group, the medium was changed to glucose-free DMEM supplemented with 5.5 mM glucose, and the cells were then maintained at 37 °C with 5% CO₂. Finally, the medium was changed to DMEM containing 10% FBS, after which the cells were cultured for 24 hours at 37 °C with 5% CO₂.

### Flow cytometry

Firstly, bEnd.3 cells were seeded at a density of 20,000 cells per well in a 12-well plate. The siRNA transfection procedure was performed on the OGD NC siRNA and OGD *Cav-1* siRNA groups following an overnight culture. Then the OGD procedure was performed on bEnd.3 cells after 30 hours. After OGD treatment, cells were incubated in DMEM medium (10% FBS) containing either Methyl-β-cyclodextrin (MβCD, 10 mM), chlorpromazine (10 μg/mL), or DMEM medium (10% FBS) for 1 hour.

The medium was then changed to DMEM medium (10% FBS), and Cy5-ADA-labelled *P. gingivali*s (MOI = 10) was added. BEnd.3 cells were cultured at 37 °C in darkness with 5% CO₂ for 2.5 hours. Then, 10 μL of Hoechst 33258 (2 mg/mL) was added, after which the cells were maintained for an additional 0.5 hours. The cells were then washed three times with saline and maintained in saline containing gentamicin (300 μg/mL) and metronidazole (200 μg/mL) for 1 hour. The cells were washed three times with saline, collected, and fixed with 4% paraformaldehyde. The cells were analysed using a flow cytometer (BD, FACSymphony A1, USA). The endocytosis index was calculated using the formula (% positive cells × MFI)/100 [37].

### Western blot (WB) analysis

The protein was extracted from the samples using RIPA buffer and quantified using a BCA kit (Millipore, #71285M, China). 4-20% SDS-polyacrylamide gradient gels were used, with vertical electrophoresis set to 200 V for 30 minutes. After that, the proteins were transferred to a polyvinylidene fluoride membrane at 400 mA for 30 minutes. The membranes were blocked and incubated with primary antibodies at 4 °C overnight. The primary and secondary antibodies used are presented in Table S4. The ChemiDoc MP system (Bio Rad, CHEMIDOC MP, USA) was used to detect protein bands, and ImageJ (NIH, 1.54f, USA) was used to quantify the protein levels.

### Immunofluorescence staining (IF) analysis

BBB organoids of day 3 were fixed with 4% paraformaldehyde. The BBB organoids were incubated with primary antibodies, followed by the secondary antibody and DAPI. After washing with PBS, the BBB organoids were placed in an 8-well chambered cover glass for imaging using the confocal fluorescence microscopy (Olympus, SpinSR, Japan). The BOs were fixed with 4% paraformaldehyde, followed by dehydration and paraffin embedding. 8 μm tissue sections were incubated with antibodies. Detailed antibody information is presented in Table S4. Confocal fluorescence microscopy (OLYMPUS, SpinSR, Japan) was used to acquire images.

### Immunohistochemistry

After fixation with 4% paraformaldehyde, paraffin sections of the BOs were prepared and incubated with anti-p-Tau and anti-Aβ_1-42_. Each section was then labelled with a secondary antibody and incubated with DAB. Detailed antibody information is presented in Table S4. The OLYMPUS VS200 system (Japan) was used to capture images of the organoids.

### ELISA analysis

The levels of IL-6, IL-1β, and TNF-α were measured in the culture medium of BOs using ELISA kits (Neobioscience, #EHC007.96, #EHC002b.96, #EHC103a.96, China). Following the kit instructions, the samples were appropriately diluted and subsequently incubated with several working solutions. After adding the stop solution to stop the reaction, the absorption intensity of the samples was measured at OD 450 nm.

### Establishment of *P. gingivalis* bacteremia in rhesus monkeys

Transient MCAO (Tmcao) was performed in two rhesus monkeys using a minimally invasive catheterization method as described previously [21, 38]. In brief, monkeys were anesthetized, and the right middle cerebral artery was blocked for 2 hours using a Guglielmi Detachable Coil. And then the Guglielmi Detachable Coil was removed from the M1 segment to recover MCA blood flow.

The BBB permeability of the two normal control monkeys and two monkeys subjected to Tmcao for ≥ 30 days was measured using computed tomography perfusion imaging (CTP). Computed tomography (United Imaging, Uct 960+, China) was used to perform CTP. The detailed settings parameters of computed tomography were the same in our previous study [21]. BBB permeability was measured using BBB-permeability surface area product (BBB-PS). The animals were anesthetized using Zoletil 50 (0.1 mL/kg, intramuscular injection). And then their CSF was collected from the cisterna magna, and *P. gingivalis* (20,000 CFU) was injected via vein. This bacterial dose was chosen to simulate the bacteremia induced by daily activities such as chewing and tooth brushing (32 CFU/mL). After 4 hours, the animals were anesthetized, and CSF was extracted again.

CSF samples (200 μL) were used to isolate DNA utilizing a TIANamp Micro DNA Kit (Tiangen, #DP316, China). Following the instructions from the manufacturer, DNA was extracted from CSF samples. A nested-PCR method was used to detect *P. gingivalis*. In brief, 25 μL of 2×Taq Plus Master Mix Ⅱ (Vazyme, #P213, China), 0.4 μL of forward primer 1, 0.4 μL reverse primer 1, 5 μL DNA extracted from CSF, and 19.2 μL H_2_O were mixed and loaded in the PCR machine. The PCR conditions are presented in Table S5. The products were diluted 10-fold. SYBR qPCR mix (Vazyme, #Q711, China) was used to perform the second round of amplification. 10 μL SYBR, 0.4 μL forward primer 2, 0.4 μL reverse primer 2, 2 μL diluted PCR products, and 7.2 μL H_2_O were mixed and placed in the CFX96 real-time PCR Detection system (Bio-Rad, CFX Duet Real-Time PCR System, USA). The qPCR conditions were presented in Table S6. A segment of the bacterial 16S rRNA gene was synthesized into a plasmid, which was then serially diluted for the preparation of standard curve samples.

### CSF proteomics analysis

Human CSF proteomics study [39] was used, and 243 observations were extracted from the study, containing 187 normal individuals and 56 AD patients. Following normalization and log2-transformation of the protein abundance data, all values were calibrated against the control group measurements in the study [39]. Partial least squares discriminant analysis (PLS-DA) was performed to compare the two groups. Differential expression analysis was performed by using Student’s t-test, following Benjamini-Hochberg correction. Differentially expressed proteins were defined by dual thresholds: adjusted *P*-value < 0.05 and absolute log2-fold change > 0.5. Kyoto encyclopedia of genes and genomes (KEGG) pathway enrichment in the differential expression proteins was performed by using an online platform for data analysis and visualization (https://www.bioinformatics.com.cn, last accessed on 10 Dec 2024). Gene set enrichment analysis (GSEA) was also performed on the platform.

### Liquid chromatography-tandem mass spectrometry (LC-MS/MS) analysis

3 days after BBB organoids establishment, the culture medium of BBB organoids was replaced with either BBB working solution alone or BBB working solution containing tetracycline (0.05 or 0.01 mg/mL, added 10 μL) or ketamine (50 nM, 200 nM, or 400 nM), respectively. After 24 h, the culture medium and BBB organoids were collected separately. The BBB organoids were added to ultrapure water and grinding beads, then homogenized using a homogenizer. The homogenate was centrifuged, and the supernatant was collected. The culture medium and the BBB organoid supernatant were extracted with an equal volume of chloroform. The organic phase was collected, concentrated, and analyzed by LC-MS/MS.

Levels of tetracycline and ketamine in the BBB organoids were quantified using LC-MS/MS. The system comprised a Waters HPLC coupled to an AB Sciex API 5000 mass spectrometer. Chromatographic separation was achieved using a Waters column (50 mm x 2.1 mm i.d.; 1.7 µm particle size) maintained at 40 °C. A mobile phase gradient (detailed in Table S7) was delivered at a flow rate of 0.60 mL/min, with an injection volume of 5 µL. Dexamethasone and verapamil served as internal standards. Data acquisition and processing were performed using Analyst^®^ software (Gerstel GmbH & Co. KG).

### Statistical analysis

All data were presented as the mean ± SEM. All statistical procedures were implemented in GraphPad Prism (v8.0.2), with one-way ANOVA applied for tri-group comparisons and two-tailed Student’s t-tests for inter-group analyses. Statistical significance was defined as *P* < 0.05.

## Results

### BBB impairment and immune dysregulation were observed in AD’s pathogenesis

CSF biomarkers exhibit high sensitivity and anatomical specificity, enabling direct characterization of intracerebral pathological features, which confers distinct advantages in neuroscience research. To elucidate the functional characteristics of the BBB and investigate the cerebral innate immune status in AD patients, we performed bioinformatics analyses on existing CSF proteomics datasets [39]. As shown in Fig. 1A, AD and control (with normal cognition and normal amyloid markers expression) demonstrate discernible segregation. We further conducted the volcano plot to visualize the proteomic difference between the AD and control groups. Of the 1091 proteins analyzed, 707 were significantly upregulated, and 384 were significantly downregulated (Fig.1B). Specifically, in the AD group, this differential expression was reflected by significantly elevated levels of blood-derived proteins (ALB, IGG-1, FGA, FGB) and downregulated expression of LRP1 and PDGFRβ—two proteins essential for maintaining normal BBB function (Fig. 1C). Notably, key components of the complement system (C4A and C4B) were also remarkablely upregulated in the AD group, suggesting brain immune activation (Fig. 1C).

**Fig. 1.**
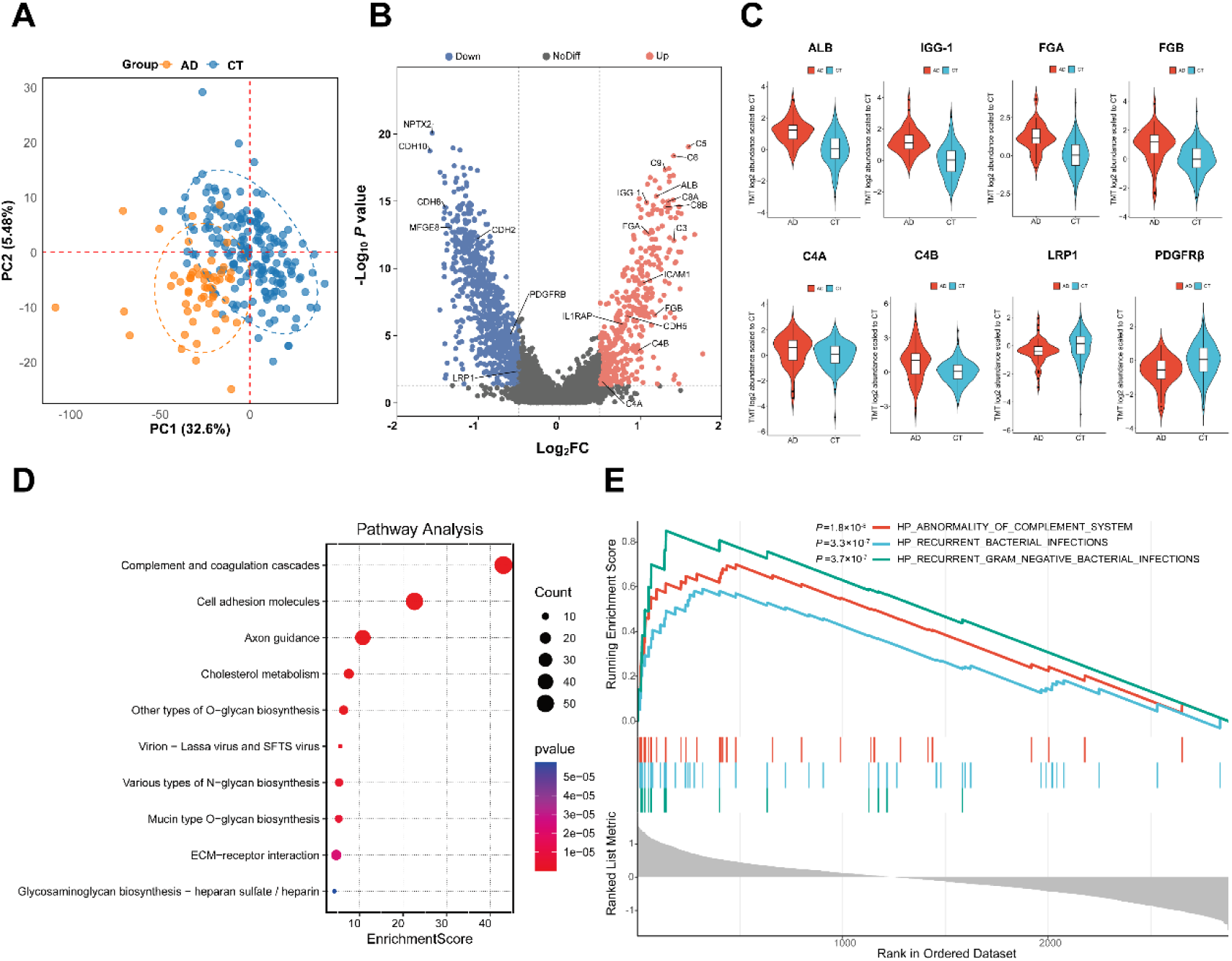
BBB impairment and immune dysregulation were observed in AD’s pathogenesis. (**A**) PLS-DA analysis of AD patients and controls. (**B**) Volcano plot of differentially expressed proteins in the CSF proteome with AD relative to control cases. (**C**) Violin and box plots of representative proteins differentially expressed in CSF between AD and controls. ALB (log_2_FC = 1.20, *P* = 4.33×10^-16^), IGG-1 (log_2_FC = 1.08, *P* = 1.24×10^-15^), FGA (log_2_FC= 1.10, *P* = 2.63×10^-13^), FGB (log_2_FC = 1.18, *P* = 2.95×10^-7^), C4A (log_2_FC = 0.53, *P* = 0.03), C4B (log_2_FC = 0.98, *P* = 1.23×10^-4^), LRP1 (log_2_FC= -0.52, P = 0.0045) and PDGFRβ (log_2_FC = -0.60, *P* = 7.33×10^-6^). (**D**) Dot plot of the KEGG enrichment analysis on the different expressed proteins in CSF with AD relative to control cases. (**E**) Enrichment plot from GSEA comparing AD to CT for different expressed proteins in CSF. n = 56 (AD), n = 187 (CT).

In the KEGG analysis, 42 significantly enriched pathways (*P* < 0.05) were identified in the AD group. Among them, the top-ranked pathway was complement and coagulation cascades (Fig. 1D). In the GSEA, abnormality of the complement system was also significantly enriched (Fig. 1E) in the AD group. In addition, recurrent bacterial infections and recurrent gram-negative bacterial infections were remarkably enriched in the AD group.

### BBB disruption facilitated *P. gingivalis* brain invasion *in vivo*

Chronic cerebral hypoperfusion plays an important role in the early pathophysiological progression of AD, but the causality between chronic cerebral hypoperfusion and neuronal damage remains controversial [40]. We constructed a rat 2VO model to simulate chronic cerebral hypoperfusion, ensuring that only vascular detrimental factors were involved [26]. As shown in Fig. 2A, the CBF decreased significantly on day 1 (64.8 ± 3.3%) and day 3 (71.9 ± 3.7%) after 2VO surgery, indicating the chronic cerebral ischemia model was successfully established. To explore the change of BBB permeability, Evans blue was used as a tracer (Fig. 2B). From day 1 to 3, compared with the SHAM group, the permeability of BBB increased significantly in both the hippocampus and cortex of the rat brain (Fig. 2C).

**Fig. 2.**
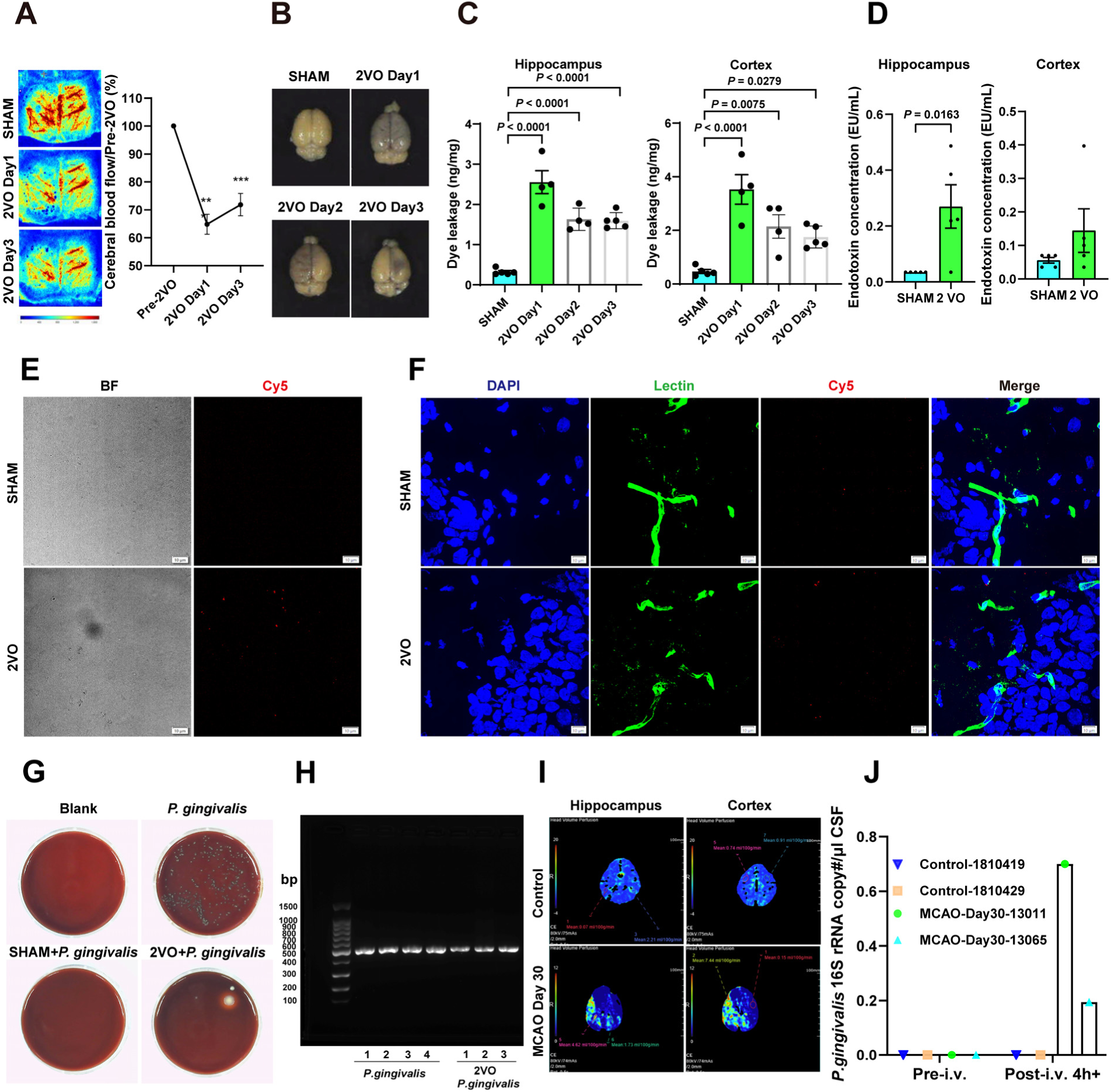
BBB disruption facilitated *P. gingivalis* brain invasion *in vivo*. (**A**) Left are the representative contrast images, right is the quantification of CBF at baseline and on day 1 and day 3 post-2VO (n = 7). ** *P* < 0.01, *** *P* < 0.001. (**B**) Evans blue staining was performed to detect BBB integrity. (**C**) Quantification of Evans blue at various time points (n = 4-5). (**D**) Endotoxin levels in the hippocampus and cortex of rats following infection with a pathophysiological dose of *P. gingivalis*, data were presented as mean ± SEM, n = 5. (**E**) Rats were infected with Cy5-ADA-labeled *P. gingivalis*, and CSF was collected and mixed with 0.5% agarose, spread on a slide, sealed with a coverslip, and then imaged. (n = 4). (**F**) IF analysis of lectin staining of the hippocampus in rats infected with Cy5-ADA-labeled *P. gingivalis*. DAPI (blue), Lectin (green), and Cy5 (red) scale bar:10 μm (n = 4). (**G**) Anaerobic culture of CSF samples from infected rats (n = 4). 60 mm diameter Petri dish. (**H**) PCR analysis of colony DNA from anaerobic cultures of *P. gingivalis* positive controls and CSF from infected 2VO rats. (**I**) Representative images of BBB-PS in control and MCAO rhesus monkeys. (**J**) PCR analysis of *P. gingivalis* in rhesus monkeys’ CSF. The data were normalised according to CSF volume (n = 2).

Oral activities such as toothbrushing and chewing may increase the prevalence of bacteremia, with blood bacterial concentrations of 0.97–32 CFU/mL [41]. Periodontal scaling procedures can introduce bacteria into the bloodstream of patients with periodontitis. A blood bacteria level of 512 CFU/mL can be caused by pre-procedural mouth care for patients with periodontitis [31]. To simulate detectable bacteremia in rats, a bacterial load of 500 CFU/mL was administered. To assess the ability of *P. gingivalis* or its products to cross the BBB when its permeability was elevated, we administered *P. gingivali*s intravenously to the rats through the tail vein at the pathophysiological concentrations (5000 CFU) [31] for 3 consecutive days, and the LPS levels in brain tissue were subsequently assessed. LPS is present on the outer membrane of *P. gingivalis*, and the colourimetric LPS assay exhibits a high sensitivity of 0.01 EU/mL [42]. As shown in Fig. 2D, LPS level in the hippocampus of the 2VO group (0.270 ± 0.078 EU/mL) was significantly higher than in the SHAM group (0.035 ± 0.000 EU/mL). In addition, the LPS in the cortex of the 2VO group showed an increasing trend (Fig. 2D). These results indicated that *P. gingivalis* or LPS had invaded the brain.

*P. gingivalis* were firstly labeled by Cy5-ADA (as shown in Fig. S1), and subsequently injected via the tail vein into the rats. Fluorescent signal was detected exclusively in the CSF of the 2VO-infected group (Fig. 2E). As shown in Fig. 2F and Fig. S2, compared with the SHAM-operated group, Cy5-fluorescent signal was detected in extravascular areas (lectin-negative regions) of the hippocampus and cortex vasculature in the 2VO-infected group. These findings collectively demonstrated that 2VO-induced increases in BBB permeability, allowing *P. gingivalis* to infiltrate the brain parenchyma, leading to bacterial colonization in both the hippocampus and cortex.

To further confirm that *P. gingivalis* has invaded the brain of 2VO rats, we extracted their CSF for anaerobic culture. As shown in Fig. 2G, no bacterial colony growth was observed in either the blank control group or the SHAM-infected group. Characteristic colonies were detected in all four plates of the *P. gingivalis* group, while colony growth was observed in one plate of the 2VO-infected group. Furthermore, DNA extracted from the colonies was subjected to PCR analysis, which showed identical results to the DNA extracted from the *P. gingivalis* (Fig. 2H). The PCR bands were excised, sequenced, and further verified to originate from *P. gingivalis* (Table S8).

To accurately simulate human vascular pathologies, we conducted experiments in rhesus monkeys subjected to MCAO modelling. First, we selected 2 male rhesus monkeys at 30 days post-MCAO surgery and two non-surgery control monkeys, then conducted CTP with an iodinated contrast agent to assess their BBB permeability. As shown in Fig. 2I, in the ipsilateral hippocampus and cortex, the BBB-permeability surface area product (BBB-PS) was higher than that in the control group. After the monkeys were administered with *P. gingivalis* intravenously at the pathophysiological concentration levels (20,000 CFU), *P. gingivalis’* DNA was detected in the MCAO group only (Fig. 2J). These data showed that the increased BBB permeability in MCAO monkeys facilitated the invasion of *P. gingivalis* into the brain. These results suggested that *P. gingivalis* at pathological physiological concentration levels more readily invades animals with compromised BBB.

### Enhanced transcytosis, not tight junction breakdown, underlies BBB disruption in 2VO rat

To further elucidate the pathways through which *P. gingivalis* crosses the BBB, we isolated rat hippocampal and cortical microvessels and extracted the RNA for RT-qPCR analysis. As shown in Fig. 3A, compared with the SHAM group, only the expression level of *Ocln* decreased remarkably after the 2VO surgery in the hippocampus, while the other two tight junction-related genes, *Tjp1* or *Cldn5*, showed no difference. In contrast to the results from the hippocampus, compared with the SHAM group, we only observed significantly decreased expression of the *Ocln* gene on day 1 after 2VO surgery in the cortex (Fig. 3B). To characterize the structural damage of the BBB following 2VO modelling, we performed TEM analysis on hippocampal and cortical microvessels of the rats. As shown in Fig. 3C and D, both the SHAM group and the 2VO group exhibited structurally intact tight junctions in the hippocampal and cortical BBB, with no gaps or disruptions observed. These results suggested that *P. gingivalis* may not infiltrate the BBB through a paracellular way.

**Fig. 3.**
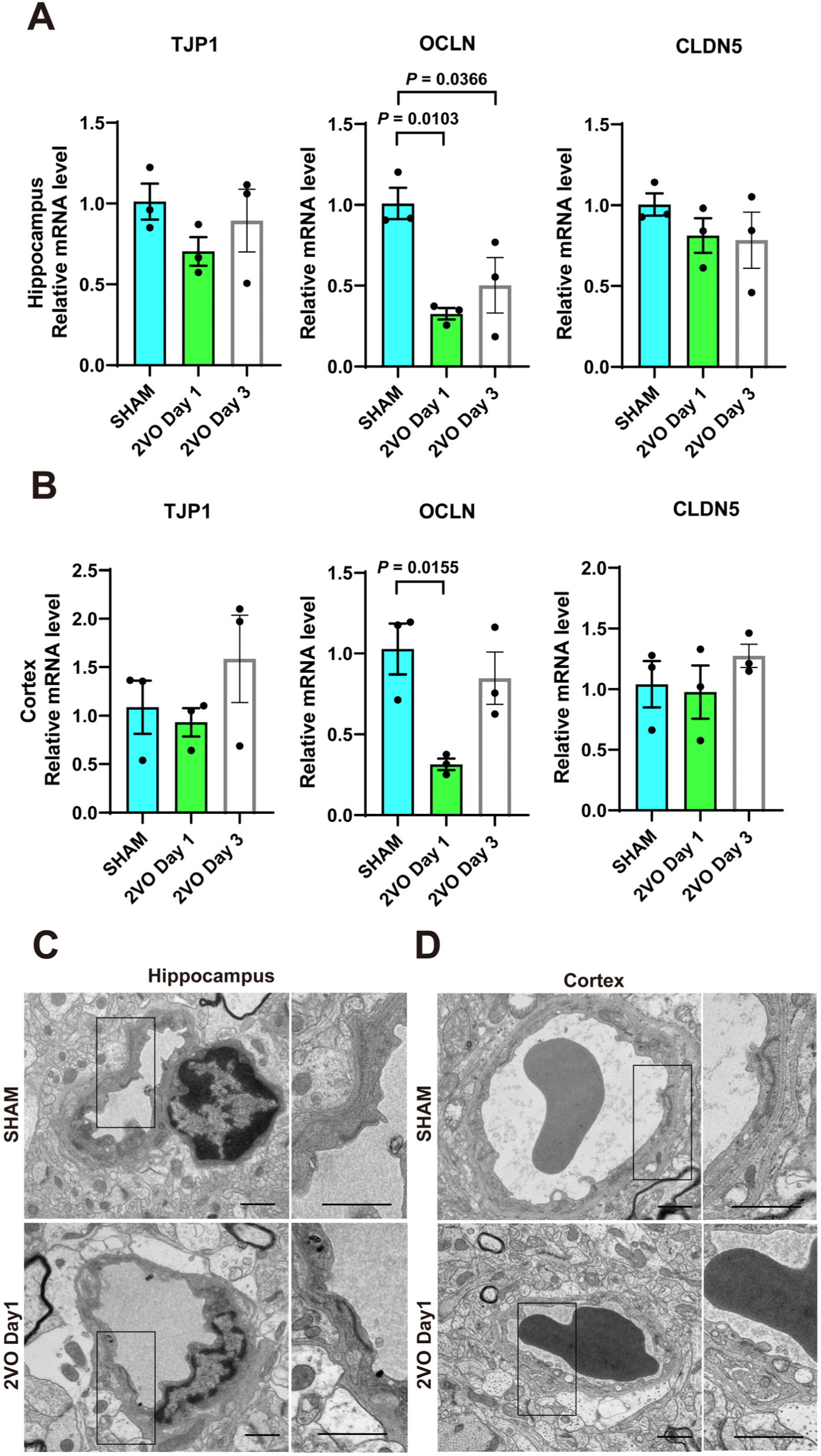
The tight junctions of the cerebral endothelial cells remained intact following 2VO surgery. (**A**, **B**) RT-qPCR analysis of genes associated with tight junction proteins (n = 3), data were presented as mean ± SEM. (**C**, **D**) TEM images of the tight junctions of hippocampal and cortical capillaries in the SHAM and 2VO Day 1 groups. Scale bar: 1 μm.

Interestingly, the 2VO group exhibited markedly increased caveolae formation (black arrows) in hippocampal and cortical (Fig. 4A) vascular endothelia. As shown in Fig. 4B, in the hippocampus, we found that the expression level of *Mfsd2a*, a key gene involved in non-specific transcytosis of the BBB, decreased significantly post-2VO (on day 1 and 3). However, we only observed that the expression level of *Mfsd2a* decreased remarkably post-2VO on day 1 in the cortex. MFSD2A regulates the permeability of the BBB by mediating the transport of ω-3 fatty acids (such as DHA) in BBB endothelial cells, thereby inhibiting caveolae formation [43, 44]. Since endothelial caveolae are primarily formed by the Cav-1 protein, we further examined the dynamic changes in Cav-1 and MFSD2A protein expression. Compared with the SHAM group, we only observed a significant decrease in the expression level of MFSD2A on day 3 (*P* = 0.0130) following 2VO surgery in the hippocampus, (Fig. 4C). In addition, the expression level of Cav-1 increased significantly on day 1 and 3 in the hippocampus, while on day 1, the expression level of Cav-1 in the cortex showed a trend of increasing. Interestingly, compared to the cortex, the hippocampus exhibited significantly higher Cav-1 expression levels both on day 1 (*P =* 0.0306) and day 3 (*P =* 0.0075). An enhanced transcytosis activity may suggest that *P. gingivalis* may infiltrate the BBB through a transcellular pathway.

**Fig. 4.**
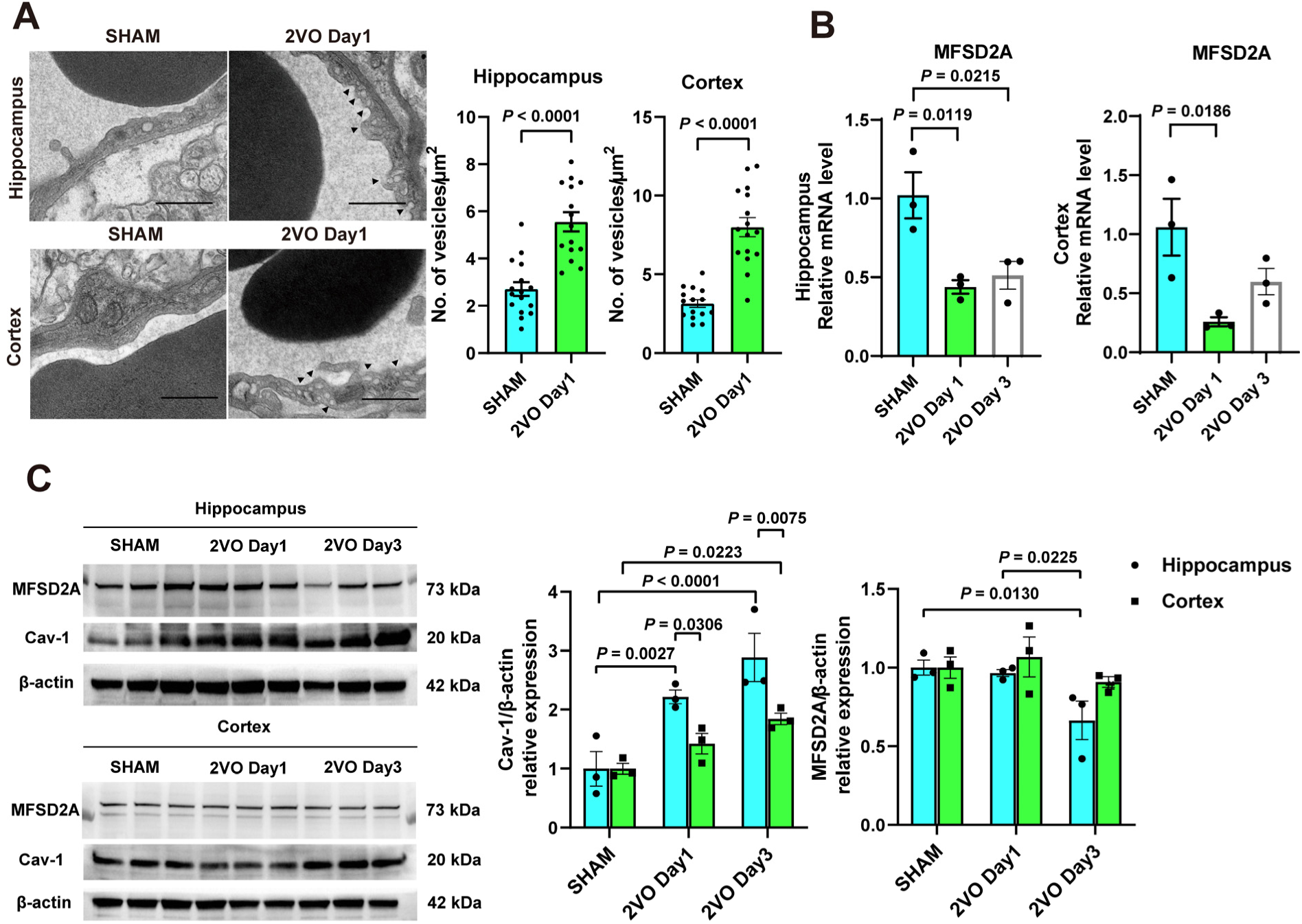
Following 2VO, the level of endothelial vesicles in cerebral vascular endothelial cells increased. (**A**) The left image showed TEM images of endothelial vesicles in the hippocampal and cortical capillaries in the SHAM and 2VO Day 1 groups. The right charts showed the quantification of the number of vesicles in endothelial cells (n = 16). Scale bar: 1 μm. (**B**) RT-qPCR analysis of genes associated with vascular function (n = 3), data were presented as mean ± SEM. (**C**) Cav-1 and MFSD2A expression in the hippocampus and cortex at different time points (n = 3). Relative protein expression levels are shown in the bar chart. Data were presented as mean ± SEM (n = 3).

### Disruption of the BBB organoids facilitated *P. gingivalis* invasion

To investigate whether increased permeability of the BBB facilitates *P. gingivalis* penetration and to explore the underlying mechanisms, we constructed self-assembled BBB organoids for *in vitro* studies. As shown in Fig. 5A, after 3 days, human cerebral microvascular endothelial cells (hCMEC/D3) cells and pericytes were localized to the peripheral margins of the BBB organoids, exhibiting high ZO-1 expression, while astrocytes populated the central core compartment. To better analyze the function of BBB organoids, we introduced two chemicals—ketamine and tetracycline—into the culture medium. Ketamine is known to penetrate the BBB [45], whereas tetracycline cannot [46]. As shown in Fig. S3, ketamine was detected in the homogenate of the BBB organoids, while tetracycline was almost absent. These results demonstrate that the BBB organoids exhibit selective permeability to compounds.

**Fig. 5.**
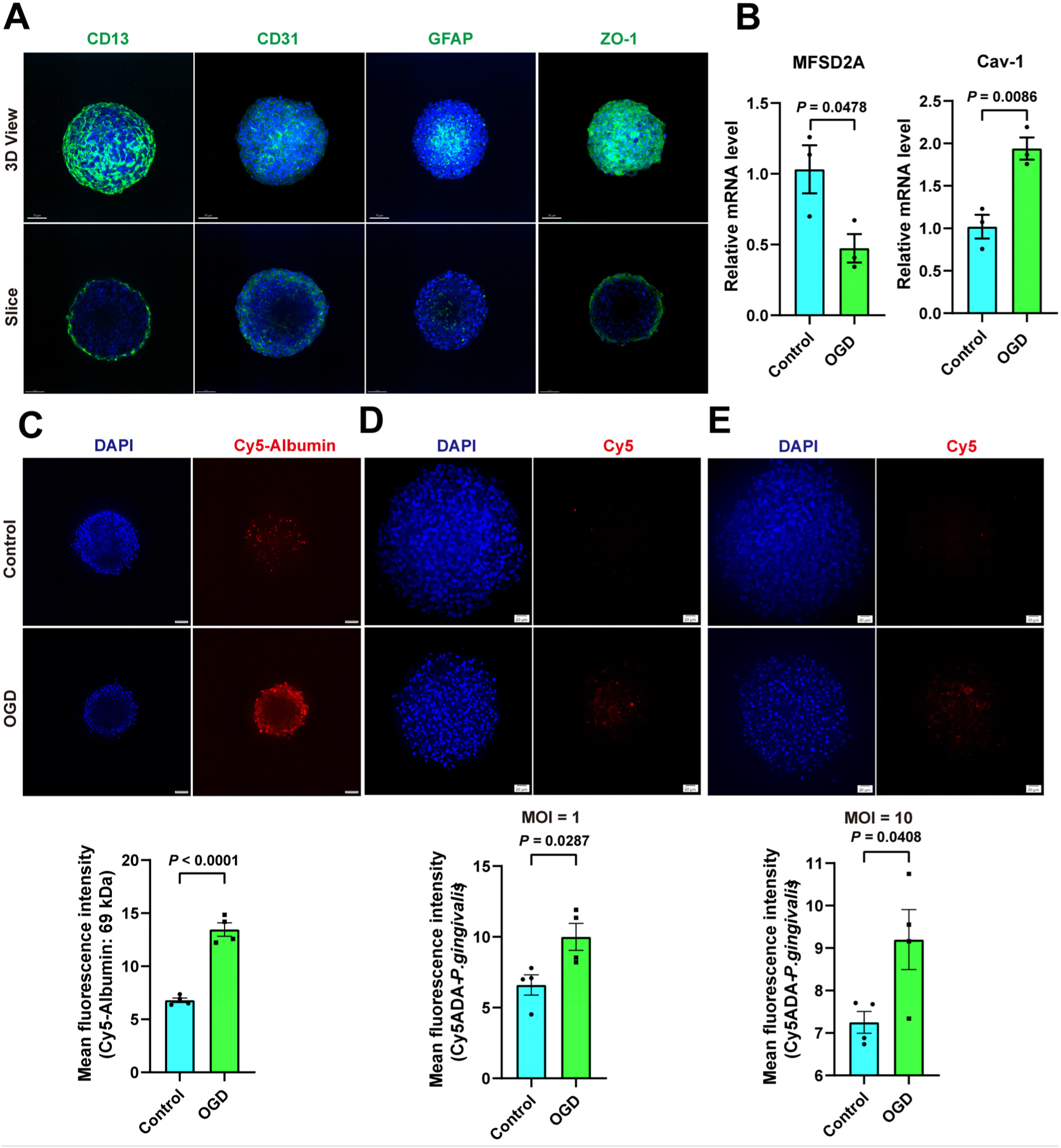
*P. gingivalis* infiltrates BBB organoids treated with OGD treatment. (**A**) IF analysis of BBB organoids, DAPI (blue), CD13, CD31, GFAP, and ZO-1 (green), scale bar: 70 μm. (**B**) RT-qPCR analysis of gene expression relative to *ACTIN* in BBB organoids. Data were presented as mean ± SEM, n = 3. (**C**) The fluorescence intensity of Cy5-albumin was quantified in BBB organoids, scale bar: 50 μm, (n = 4). (**D**, **E**) IF analysis of BBB organoids following infection with Cy5-ADA-labeled *P. gingivalis*. The top panel shows representative IF images, and the bottom panel shows the quantification of the Cy5 fluorescence intensity in BBB organoids. Data were presented as mean ± SEM, scale bar: 20 μm, n = 4.

To model the ischemic condition of the BBB, we subjected the BBB organoids to OGD treatment. The expression level of *MFSD2A* in the BBB organoid was reduced, while the expression level of *CAV-1* was increased significantly in the OGD group (Fig. 5B), consistent with our observations in the *in vivo* experiments, suggesting that the transcellular pathway of the BBB was activated. We further used Cy5-albumin to evaluate BBB permeability. Compared to the control, we found that their permeability increased significantly in the OGD group (Fig. 5C). OGD-induced BBB disruption significantly potentiated *P. gingivalis* invasion across this barrier, as evidenced by elevated fluorescence signals in treated organoids at both MOI 1 (Fig. 5D) and MOI 10 (Fig. 5E) compared to the control.

### Post-OGD bEnd.3 cells internalized *P. gingivalis* via Cav1-mediated endocytosis

According to UniProt, the amino acid sequences of human Cav-1 share high similarity with those of mouse and rat Cav-1, showing 94.94% and 94.38% identity, respectively, indicating only minor differences among species. Cav-1 is expressed in cerebral vascular endothelial cells across humans, mice, and rats. Furthermore, the mouse brain microvascular endothelial cell line bEnd.3 has been extensively utilized to study bacterial and nanomaterial transcytosis across the BBB [14, 47, 48]. To investigate the mechanism by which *P. gingivalis* takes advantage of elevated BBB permeability to cross the BBB, we conducted more downstream experiments using bEnd.3 cells. First, we examined the changes in Cav-1 expression following OGD treatment. Compared to the control, the OGD-treated group exhibited a significant increase in Cav-1 expression (Fig. 6A).

**Fig. 6.**
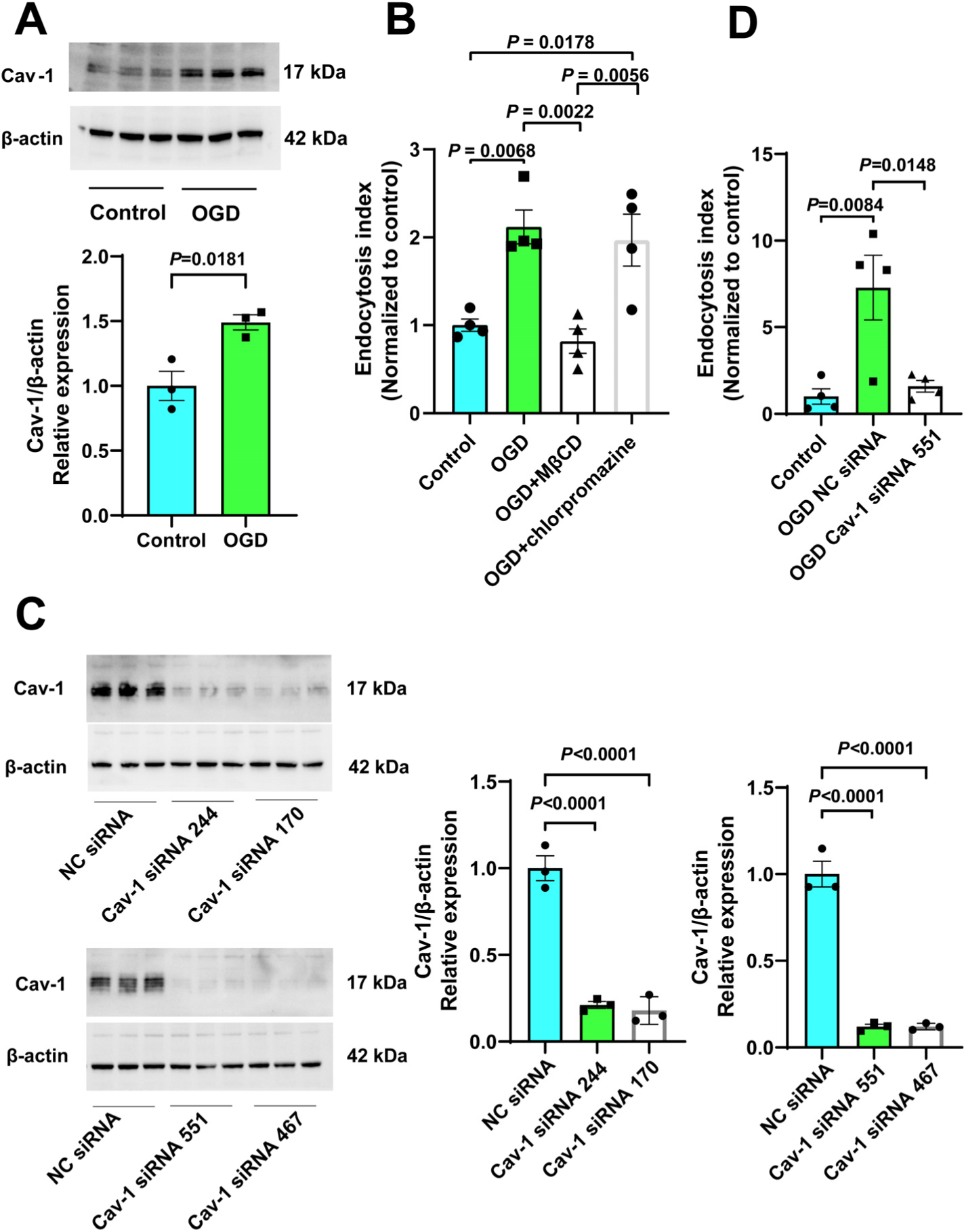
Post-OGD bEnd.3 Cells internalized *P. gingivalis* via Cav-1-mediated endocytosis. (**A**) WB analysis of control and OGD, with β-actin used as an internal standard in mouse bEnd.3 cells. Data were presented as mean ± SEM, n = 3. (**B**) Flow cytometric detection of *P. gingivalis* (labelled with CY5-ADA, MOI = 10) in bEnd.3 cells. The image shows the endocytosis index among the different groups. Data were presented as mean ± SEM, n = 4. (**C**) WB analysis was performed to detect the level of Cav-1 in bEnd.3 cells treated with *Cav-1* siRNA or NC siRNA. Quantification of relative expression was conducted and data were presented as mean ± SEM, n =3. (**D**) Flow cytometry was used to detect *P. gingivalis* in bEnd.3 cells in the control, OGD NC siRNA, and OGD *Cav-1* siRNA groups. Data were presented as mean ± SEM, n = 4.

Subsequently, bEnd.3 cells were co-cultured with Cy5-ADA-labeled *P. gingivalis* at low MOI (10), and the endocytic index was quantified via flow cytometry using a gating strategy as illustrated in Fig. S4. Compared to the control, OGD-treated bEnd.3 cells exhibited significantly enhanced internalization of *P. gingivalis.* MβCD, a caveolae-disrupting agent, suppressed *P. gingivalis* internalization in OGD-treated bEnd.3 cells, whereas chlorpromazine, an inhibitor of clathrin-mediated endocytosis, showed no effect, suggesting that OGD potentiates bacterial invasion primarily via Cav-1-dependent pathways (Fig. 6B). Furthermore, Cav-1 knockdown via four independent *Cav-1* siRNAs reduced Cav-1 protein levels significantly compared to negative control (NC) siRNA (Fig. 6C). OGD enhanced *P. gingivalis* internalization in NC siRNA-treated cells, an effect that was eliminated by *Cav-1* siRNA_551 treatment (Fig. 6D). These results indicate that OGD-induced Cav-1 upregulation promotes bacterial transcytosis by increasing caveolae-mediated transcellular permeability.

### *P. gingivalis* induced AD-like pathophysiological features in human brain organoids

BOs exhibit no species differences that can accurately reflect the molecular mechanisms of human diseases, offering multiple significant advantages in neuroscience research [34, 49]. Therefore, we leveraged human BO to test the hypothesis that *P. gingivalis* infection triggers AD-associated pathological cascades. To assess glial maturation dynamics, RT-qPCR was performed on BOs at 66 and 94 days. Progressive upregulation of microglial markers and astrocytic *GFAP* was observed in mature BOs (Fig. 7A). To further confirm that BOs contain microglia, we performed RNA-Seq, and as shown in Fig. S5, microglia were detected.

**Fig. 7.**
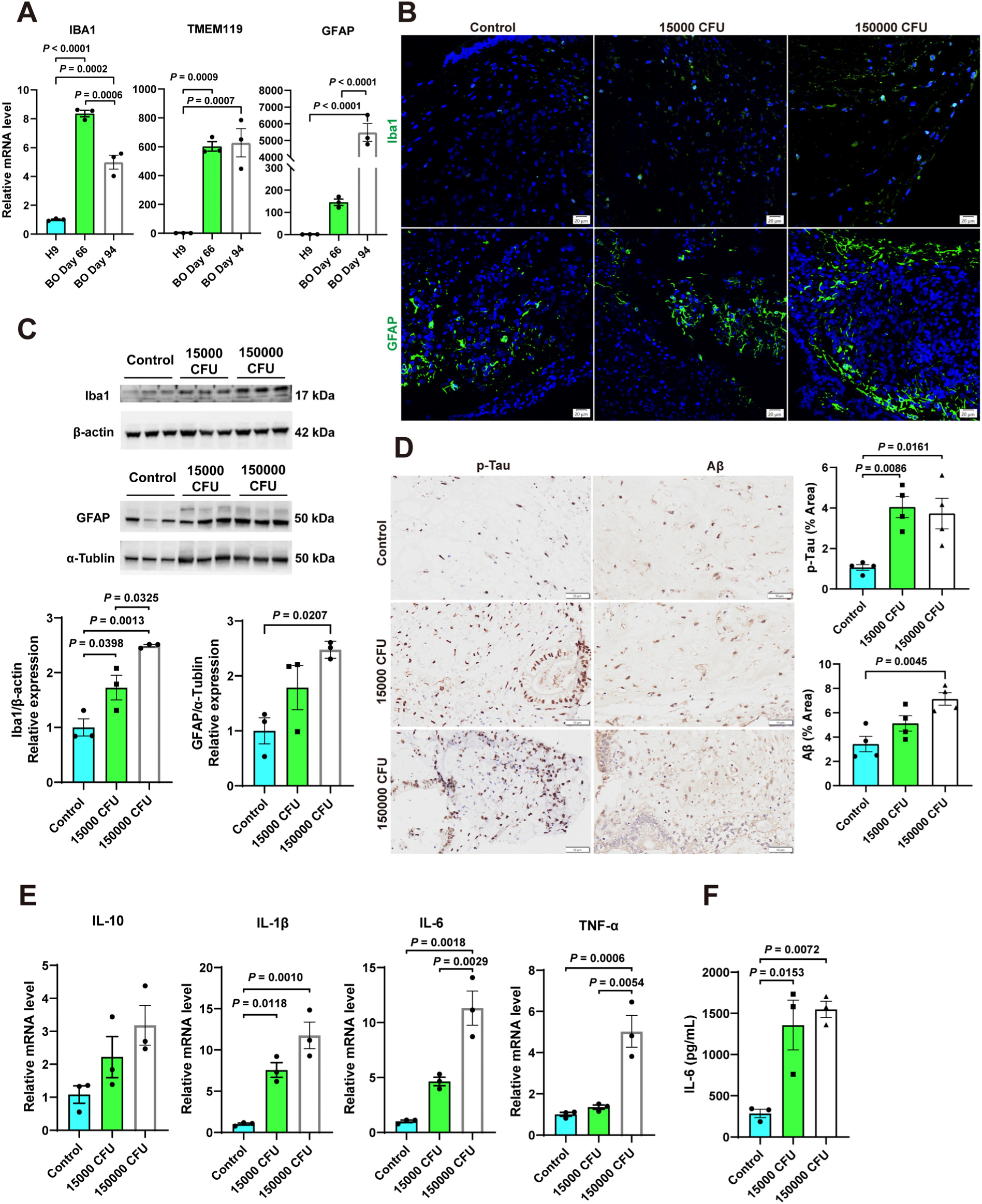
*P. gingivalis* induced AD-like pathophysiological features in human BOs. (**A**) RT-qPCR analysis of gene expression relative to *ACTIN* in BOs and H9 ESCs. Data were presented as mean ± SEM, n = 3. (**B**) Representative images of control, 15,000 CFU, and 150,000 CFU sections of BOs sections stained with anti-Iba1 or anti-GFAP antibodies. Scar bar: 20 μm, n = 5. (**C**) WB analysis of control, 15,000 CFU, and 150,000 CFU in BOs, with β-actin or α-Tublin as an internal standard. Data were presented as mean ± SEM, n = 3. (**D**) The expression of p-Tau and Aβ in BOs following co-culture with *P. gingivalis* or not. Scale bar: 50 μm, data were presented as mean ± SEM, n = 4. (**E**) RT-qPCR analysis of gene expression in BOs following infection with *P. gingivalis* or not. Data were presented as mean ± SEM, n = 3. (**F**) The level of IL-6 in the culture medium of BOs was quantified by ELISA. Data were presented as mean ± SEM, n = 3.

Exposure to *P. gingivalis* triggered glial activation, as evidenced by elevated GFAP (astrocytes) and Iba1 (microglia) expressions in the infected BOs (Fig. 7B). WB revealed dose-dependent neuropathological changes: Iba1 levels were significantly elevated in both low and high-dose groups versus controls. GFAP expression increased only in the high-dose treatment, with no significant change in the low-dose group (Fig. 7C). Co-culture with *P. gingivalis* induced dose-dependent neuropathology in the BOs: p-Tau deposition was elevated in both low (15,000 CFU) and high-dose (150,000 CFU) groups versus controls (Fig. 7D), while Aβ accumulation was observed only in the high-dose group (Fig. 7D). Subsequent RT-qPCR analysis revealed pronounced neuroinflammation, with upregulation of *IL-6*, *IL-1β*, and *TNF-α* in the high-dose group *VS.* controls (Fig. 7E), implicating glial-driven cytokine storms in AD-like pathogenesis.

To further validate protein-level changes in inflammatory cytokine expression, we analyzed the culture medium of BOs for IL-1β, IL-6, and TNF-α. The control group exhibited an IL-6 concentration of 286.2 ± 49.69 pg/mL, while the low- and high-dose infection groups showed increased IL-6 levels of 1358 ± 302.0 pg/mL and 1547 ± 99.75 pg/mL, respectively (Fig. 7F). However, the absorbance values of IL-1β and TNF-α in all experimental groups were below the lower limit of quantification of the enzyme-linked immunosorbent assay (ELISA) kits (IL-1β < 7.8 pg/mL; TNF-α < 15.6 pg/mL). Collectively, these results indicated that co-culturing with *P. gingivalis* triggered the neuroinflammation in BOs.

## Discussion

Accumulating studies support the pathogen hypothesis of AD pathogenesis. Pathogens (e.g., HSV-1, *Chlamydia pneumoniae*, *Burkholderia sp,* and *P. gingivalis*) may be involved in the pathogenesis of AD by inducing Aβ deposition and neuroinflammation. For example, the DNA of *P. gingivalis* was detected in the CSF of AD patients [4], and *P. gingivalis* has been reported to induce AD-like pathophysiological features *in vitro* and *in vivo* [8, 50–54]. However, key gaps remain in the pathogen hypothesis of AD pathogenesis. Under normal conditions, the BBB tightly regulates the exchange of substances between blood and brain, blocking blood-borne toxins such as pathogens from entering the brain, and thereby preserving intracerebral homeostasis. Pathogens are unlikely to be able to open the BBB at the pathophysiological level (ie, blood bacteria concentration is usually at 0.97-32 CFU/mL following oral daily activities) [41]. However, increased BBB permeability has been reported as an early event in AD [55], particularly in the hippocampus, and often precedes the onset of cognitive impairment. We therefore propose that increased BBB permeability triggered by cerebrovascular pathologies during AD progression may provide a ‘window of opportunity’ for brain invasion by pathogens such as *P. gingivalis*. To test this hypothesis, we conducted a series of experiments to investigate whether *P. gingivalis* exploits enhanced BBB permeability (e.g., induced by cerebral hypoperfusion) to translocate into the brain. We further elucidated the specific pathways mediating this translocation and determined whether brain invasion elicits AD-like pathophysiological features.

Brain capillary damage and BBB breakdown in the hippocampus occur early in cognitive decline, and precede any changes in classic Alzheimer’s biomarkers like Aβ and tau [55]. Our analysis of CSF differential proteins, corroborated by evidence of perivascular blood-derived protein deposition in over 20 AD post-mortem studies [56], provided definitive support for the existence of pathological BBB leakage in AD patients. Concurrently, levels of the pericyte marker PDGFRβ were significantly reduced in the AD group, aligning with observed pericyte loss in AD animal models and post-mortem studies [57, 58]. Furthermore, LRP1 levels were significantly diminished in the AD group, potentially impairing Aβ clearance capacity within the brain [59]. In the KEGG enrichment analysis, differentially expressed proteins were significantly enriched in the complement system activation pathway, consistent with previous reports [60]. Notably, complement activation – a core component of the innate immune system – is well-documented in post-mortem brain tissue and CSF from individuals with AD [61, 62]. It is established that both Aβ and NFTs can activate the complement system [63, 64]. Complement system activation is strongly implicated in AD pathology and is a hallmark of chronic infections (e.g., periodontitis) [65]. *P. gingivalis* activates complement independently of Aβ [66], suggesting that pathogen infection may represent an additional driver of complement dysregulation in AD. Furthermore, GSEA analysis revealed significant enrichment of pathways associated with recurrent gram-negative bacterial infections in the AD group. Supporting the relevance of this finding, prior studies have detected LPS – a cell wall component of gram-negative bacteria – in the perivascular spaces of AD patients [20]. These observations collectively suggested that pathogens such as *P. gingivalis* may exploit a compromised BBB to invade the brain. Notably, Stephen S. Dominy et al. (2019) have detected *P. gingivalis* DNA in the CSF of AD patients [4]. These results suggested a possible association between BBB dysfunction in AD patients and invasion of bacteria or their products into the brain through the cerebral vasculature.

To verify that the breakdown of the BBB is a prerequisite for pathogens to enter the brain tissue, we established a 2VO model in rats without any genetic predisposition of AD. Consistent with early reports [26, 27], there was a significant decrease in CBF and a remarkable increase in BBB permeability on days 1 and 3 post-2VO surgery, compared to pre-2VO levels or SHAM. The results clearly showed that pathogens could enter the brain tissue of 2VO rats but not that of SHAM rats, indicating that the breakdown of the BBB is a prerequisite for pathogens to invade the brain tissue. The same results were also confirmed in MCAO monkeys.

Bacteremia occurs after oral procedures, such as chewing, toothbrushing, and scaling. Individuals with periodontitis have an increased risk of bacteremia during daily activities such as chewing and brushing teeth [67]. Typically, toothbrushing or chewing-induced bacteremia is of low intensity, with blood bacterial concentrations of 0.97–32 CFU/mL [41]. Rodrigo Dalla Pria Balejo et al. (2017) reported that scaling in periodontal patients leads to bacteremia, with *P. gingivalis* levels in the blood reaching 512 CFU/mL [31]. Various studies have induced bacteremia by injecting high concentrations of *P. gingivalis* into the tail vein or intraperitoneally [10, 14, 68–71]. However, bacterial concentrations in actual patient blood are lower. To simulate human pathophysiological concentration levels of bacteremia in rats at a detectable level, we used the dosage of 500 CFU/mL. To directly observe whether *P. gingivalis* at a pathophysiological dose could invade the brain following BBB disruption, we administered the bacteria via tail vein injection rather than using a periodontitis model (e.g., ligature-induced). While a periodontitis model may induce bacteremia that more closely mimics the natural disease course, the occurrence and level of bacteremia in such models are difficult to control precisely. In contrast, intravenous injection allows for the accurate delivery of a defined bacterial dose at a specific time point. We found that the LPS level increased significantly, and *P. gingivalis* was isolated from the CSF in rat with BBB disruption. Similarly, *P. gingivalis* DNA was detected only in the CSF of rhesus monkeys with an elevated BBB permeability. Other study demonstrate that long-term infection of rats with *P. gingivalis* at doses far exceeding pathophysiological concentration levels disrupts the BBB, enabling brain invasion [14]. However, the exceptionally high bacterial loads employed in such experiments drastically surpass realistic infection exposure. Thus, the general relevance of this pathological mechanism requires cautious interpretation.

In the 2VO rats, although we observed downregulation of *Ocln*, the other two crucial components of the BBB—*Tjp1* and *Cldn5*—remained relatively normal. Zhengyu Sun et al. (2021) reported that the OCLN protein level decreased on day 3 post-2VO, while the ZO-1 and CLDN5 protein levels showed no change from day 1 to day 28 post-2VO [26]. In agreement with MCAO mice and 2VO rats reports [26, 72, 73], electron microscopy in our study also revealed a relatively intact tight junction structure in both the 2VO and SHAM groups. Therefore, it is unlikely that pathogens can cross the BBB via the paracellular pathway.

Following 2VO surgery, microvascular endothelial cells in both the hippocampus and cortex exhibited increased caveolae formation, accompanied by reduced *Mfsd2a* mRNA expression and elevated Cav-1 levels, indicating an enhancement of nonspecific transcytosis. Most importantly, *MFSD2A* expression was also significantly downregulated in BBB endothelial cells from individuals with AD [74]. MFSD2A maintains a specialized lipid environment in cerebral endothelial cells that suppresses caveolae-mediated transcytosis [75]. These changes suggest that *P. gingivalis* may traverse the BBB primarily through a transcellular route.

Interestingly, in the 2VO rats, we observed that the hippocampus had a higher Cav-1 expression than the cortex, which coincided with the observation of hippocampal BBB hyperpermeability in gastrointestinal flora-disordered mice and MCAO rhesus monkeys [21, 29], suggesting that the hippocampal microvasculature may be more sensitive to ischemia, consistent with a previous report [76]. The differential expression of Cav-1 in the hippocampus and cortex likely represents a core vulnerability in the hippocampus. However, further experiments are needed to demonstrate the role of Cav-1-dependent endocytosis in neurodegenerative diseases.

Notably, BBB organoids displayed a significant increase in BBB permeability after OGD. Following co-culture with *P. gingivalis*, the bacterial fluorescence signal intensity of BBB organoids treated with OGD increased. These results indicated that increased BBB permeability directly promotes *P. gingivalis* to break through the barrier and invade the brain. To further validate the role of Cav-1 in this process, we are currently overexpressing Cav-1 in BBB organoids to assess whether Cav-1–dependent pathways enable *P. gingivalis* to cross the BBB. Using bEnd.3 cells subjected to OGD to model ischemia, we observed significant upregulation of Cav-1 expression. This finding is corroborated by our *in vivo* and organoid studies, as well as reports from other groups [77–79]. Bacterial co-culture experiments revealed significantly enhanced endocytosis of bacteria in OGD-treated bEnd.3 cells. This effect was reversed by MβCD and *Cav-1* siRNA treatment. These results demonstrated that Cav-1-dependent non-specific transcytosis constitutes the essential pathway for *P. gingivalis* translocation across the BBB. A limitation of this study is the lack of in vivo validation using Cav-1 knockout animals. Future studies will employ this model to address this gap. Previous studies have shown that when *P. gingivalis* is co-cultured with bEnd.3 cells and U-87 cells, it cannot translocate across the endothelial monolayer at an MOI of 10, and significant bacterial traversal is observed only at higher MOIs (e.g., 100 and 500) [14]. Furthermore, the upregulation of Cav-1 by *P. gingivalis* was also demonstrated at an MOI of 100 [14]. In contrast, in our BBB organoid model, following OGD treatment, *P. gingivalis* was able to cross the barrier even at an MOI as low as 1. This discrepancy suggests that a prerequisite condition—such as hypoxia-induced damage—may be necessary to upregulate Cav-1 and compromise BBB integrity, thereby enabling *P. gingivalis* translocation in a more physiologically relevant context. Although caveolae have a diameter of approximately 50 nm [80] and several studies have proposed that bacteria utilize them to cross cells [14, 81], the detailed mechanism underlying this process remains unknown.

In our study, we established human BOs and identified the expressions of microglial markers (*IBA1* and *TMEM119*), consistent with the report from Martina et al. (2024) [82], further supporting the presence of microglia in our BOs. Unlike the directed differentiation approaches, non-directed differentiation may lead to the differentiation of certain embryonic stem cells towards mesoderm, which can further differentiate into microglia [83]. Following co-culture of BOs with *P. gingivalis,* we observed increases in the positive area for both p-Tau and Aβ to different extents. Multiple *in vitro* and *in vivo* studies have demonstrated that *P. gingivalis* induces abnormal tau phosphorylation [50, 54, 84]. This aberrant phosphorylation leads to tau dysfunction and represents a key driver of AD pathogenesis. Following the *P. gingivalis* infection, we observed increased Aβ expression. This finding aligns with results from long-term infected rats [85], HSV-1-infected human BOs [86], and APP/PS1 mouse models [49], collectively suggesting that *P. gingivalis* induces Aβ deposition—a characteristic pathological feature of AD. In addition, exposure of human BOs to human serum can also induce AD-like characteristics [49, 87], suggesting that breakdown of the BBB may be a key driver in the development of AD.

Neuroinflammation mediated by activated microglia and astrocytes is closely associated with the development of AD [88]. Both our IF and WB analyses demonstrated increased expression levels of Iba-1 and GFAP in BOs following bacterial infection. Furthermore, compared to the control, the high-dose infection group exhibited upregulation of pro-inflammatory cytokines such as *IL-1β*, *IL-6*, and *TNF-α*. This increase mirrors elevated levels of IL-1β, IL-6, and TNF-α observed in brain tissue of AD mouse models [89], HSV-1-infected human BOs [86], and the cortex of C57BL/6 mice subjected to oral gavage with *P. gingivalis* LPS for 7 consecutive days [90]. Elevated levels of IL-1β, IL-6, and TNF-α in the brain are recognized markers of neuroinflammation. Activated microglia influence surrounding brain tissue by secreting pro-inflammatory cytokines such as IL-1β, IL-6, and TNF-α [91]. Our ELISA analysis confirmed significantly increased IL-6 expression in BOs following co-culture with *P. gingivalis*, which mirrored the changes observed in C57BL/6 mice exposed to *P. gingivalis* LPS *in vivo* [90]. However, the concentrations of IL-1β and TNF-α in the culture medium were below the detection limit. This is likely attributable to their inherently low levels in the medium. Studies indicate that both cortical vascular organoids and human BO-on-a-chip models cultured under homeostatic conditions typically exhibit IL-1β and TNF-α concentrations below 10 pg/mL in the culture medium [92, 93]. Collectively, these findings demonstrate that co-culture of human BOs with *P. gingivalis* promotes the activation of microglia and astrocytes, leading to the secretion of the pro-inflammatory cytokine IL-6 and the induction of neuroinflammation. This process reproduces key pathological features of AD.

## Abbreviations

2VO: Bilateral common carotid artery occlusion
AD: Alzheimer’s disease
APOE4: Apolipoprotein E4
Aβ: Amyloid-β protein
BBB: Blood-brain barrier
BBB-PS: BBB-permeability surface area product
BHI: Brain-heart infusion
BOs: Brain organoids
CAA: Cerebral amyloid angiopathy
Cav-1: Caveolin-1
CBF: Cerebral blood flow
CFU: Colony-forming unit
CSF: Cerebrospinal fluid
CTP: Computed tomography perfusion imaging
ESCs: Human embryonic stem cells
GSEA: Gene set enrichment analysis
HSV-1: Herpes simplex virus-1
IF: Immunofluorescence staining
KEGG: Kyoto encyclopedia of genes and genomes
LC-MS/MS: Liquid chromatography-tandem mass spectrometry
LPS: Lipopolysaccharide
MCAO: Middle cerebral artery occlusion
MCI: Mild cognitive impairment
MOI: Multiplicity of infection
MβCD: Methyl-β-cyclodextrin
NFTs: Neurofibrillary tangles
OGD: Oxygen-glucose deprivation
*P. gingivalis*: Porphyromonas gingivalis
PLS-DA: Partial least squares discriminant analysis
RNA-seq: RNA sequencing
TEM: Transmission electron microscopy
tMCAO: Transient MCAO
VD: Vascular dementia
WB: Western blot

## Declarations

### Ethics approval and consent to participate

All experimental protocols have been approved by the Animal Care Committee of Sichuan University West China Hospital (approval number 2020119A).

## Acknowledgments

We acknowledge the foundational work of Tijms et al. (Nat. Aging, 2024) for their AD subtype data. Mass spectrometry-based proteomic analyses were performed by the Proteomics Unit at the University of Bergen (PROBE). This facility is a member of the National Network of Advanced Proteomics Infrastructure (NAPl), which is funded by the Research Council of Norway (INFRASTRUKTUR-program project number: 295910). The authors would like to thank Sichuan Junhui Biotechnology Co., Ltd. and Singapore HumanSim Biotech Pte. Ltd. for providing the experimental platforms for this study. The authors acknowledge the technical support provided by the following staff from the Research Core Facility of West China Hospital, Sichuan University: Yi Zhang, Yue Li, Xiaojiao Wang, Dan Luo, and Xin Luo.

## Funding

Joint Funds of the National Natural Scientific Foundation of China U24A20690 Guizhou Provincial Basic Research Program (Natural Science) [2025] No. 328

Sichuan Science and Technology Program (2025ZNSFSC0703)

National Natural Scientific Foundation of China 82071349

## Authors’ contributions

Conceptualization: ZZH

Methodology: MCH, ZQX, LXJ

Investigation: MCH, ZQX, LXJ, HY, ZT, ZBC, LJS

Visualization: MCH

Supervision: ZZH

Writing—original draft: MCH, ZQX

Writing—review & editing: ZZH, ZQX

## Data availability

All data are available in the main text or the supplementary materials. Additional data related to this paper may be requested from the authors. All mass spectrometry data generated for this study with accompanying demographical information are available through the ADDI workbench (https://fair.addi.addatainitiative.org/#/data/datasets/five_csf_proteomic_subtypes_in_ad; https://doi.org/10.58085/HR6S-2991).

## Supplementary Materials

Supplementary Material 1

