## Supplementary Material 1 for "Caveolin-1–Mediated Blood-brain Barrier Transcytosis Promotes *Porphyromonas gingivalis* Invasion and Alzheimer’s Disease-Like Changes"

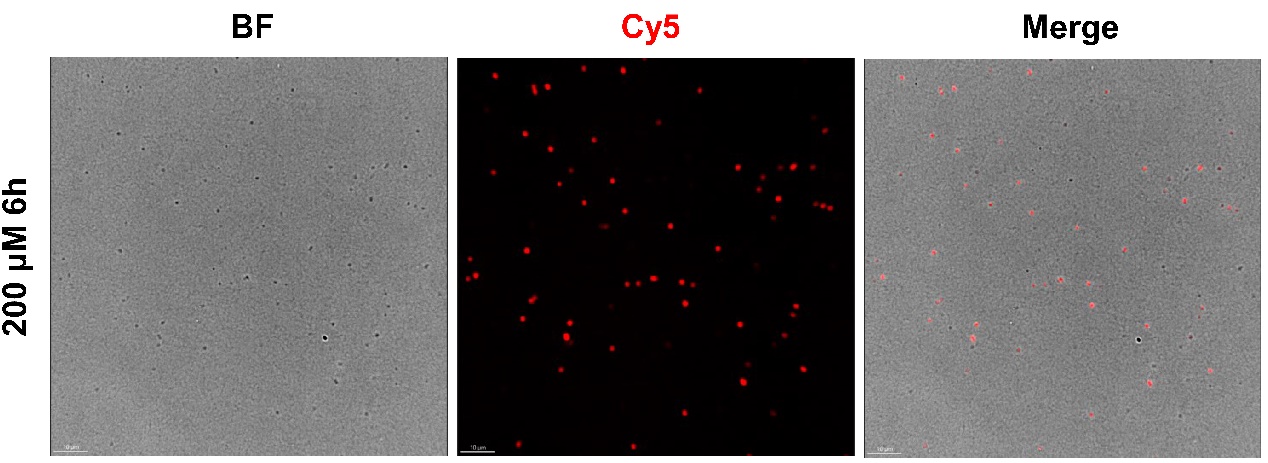

Fig. S1. *P. gingivalis* was labelled with Cy5-ADA. Cy5-ADA (red) was added to the BHI medium and cultured with *P. gingivalis* for 6 hours for bacterial labeling. Scale bar: 10 μm.

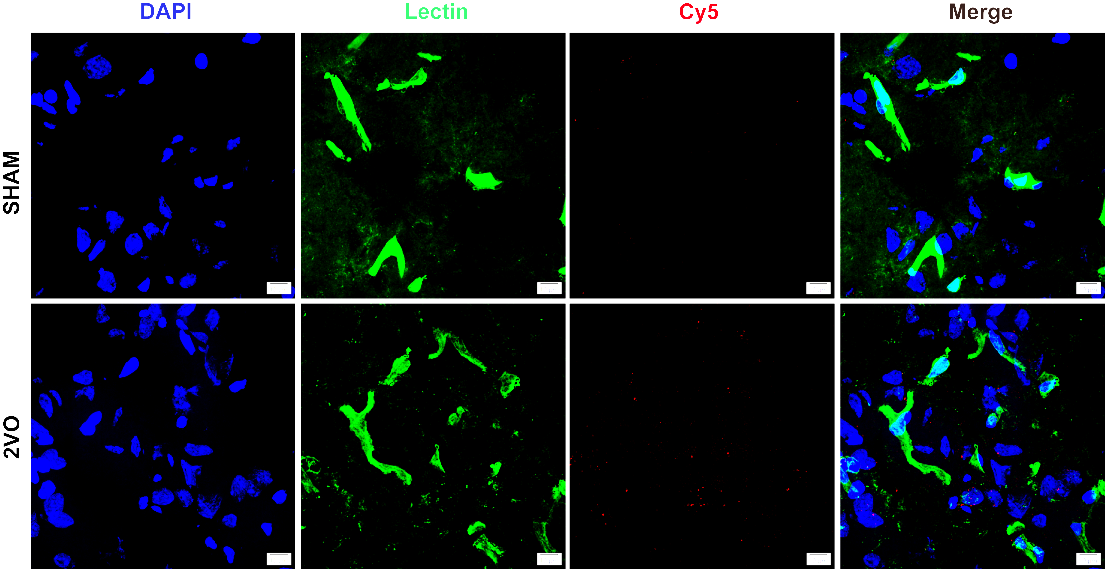

Fig. S2. Lectin staining of rat cortex. *P. gingivalis* labeled with Cy5-ADA was intravenously administered via the tail vein to rats for 3 consecutive days. Brain tissues were collected on day 4, snap-frozen, and cryosectioned for lectin staining, DAPI (blue), Lectin (green) and Cy5 (red). n = 4; Scale bar: 10 μm.

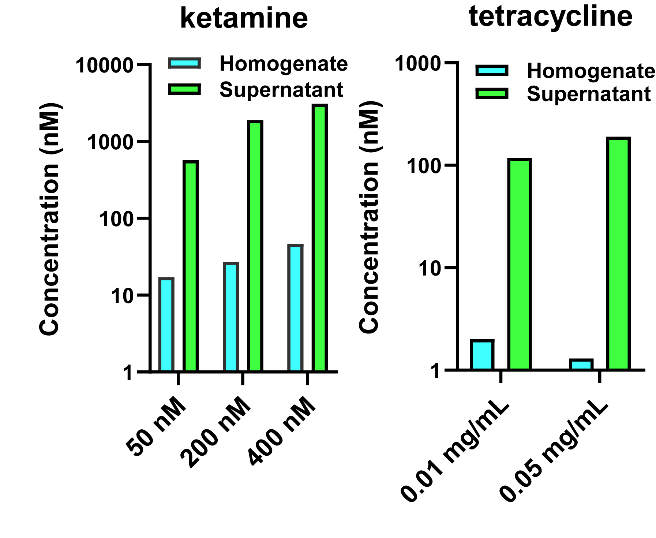

**Fig. S3. Analysis by LC-MS/MS of tetracycline and ketamine permeability using the BBB organoids.** The concentrations of tetracycline and ketamine in the homogenate and supernatant of the BBB organoids measured by LC-MS/MS.

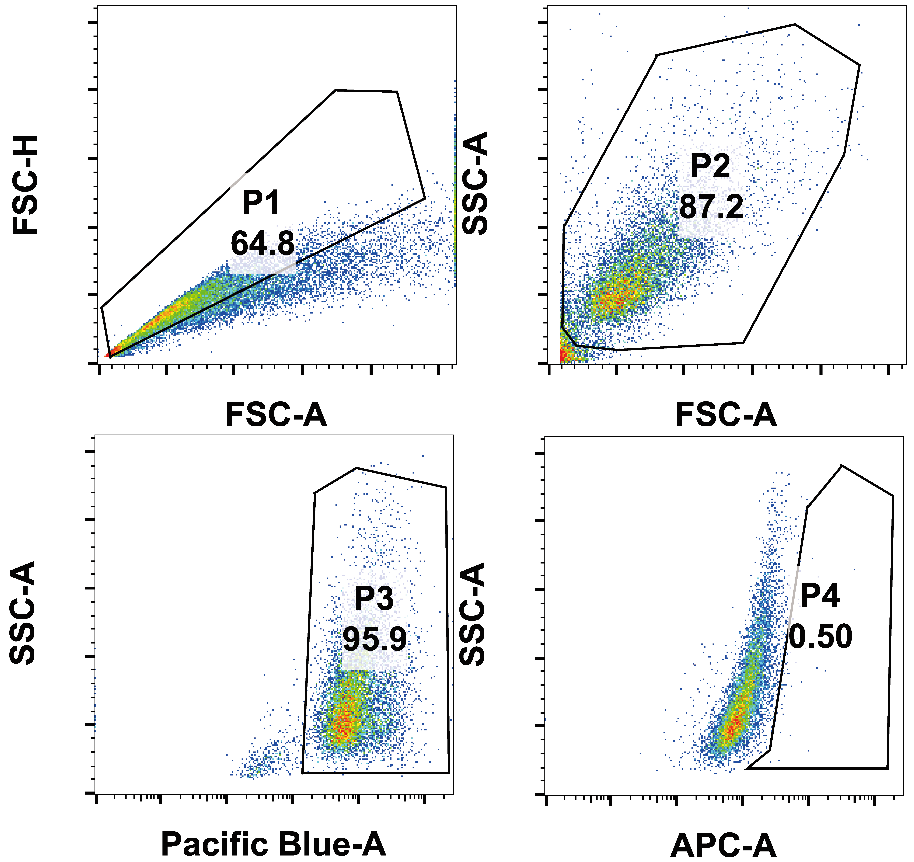

Fig. S4. Representative flow plot of gating strategy. Initially, single bEnd.3 cells are selected using the FSC-A vs. FSC-H plot (P1). Debris is excluded based on the morphological profile in the FSC-A vs. SSC-A plot (P2). Next, Hoechst 33258-stained cells are selected using the Pacific Blue-A vs. SSC-A plot (P3), and then bEnd.3 cells containing the CY5 signal are selected using the APC-A vs. SSC-A plot (P4).

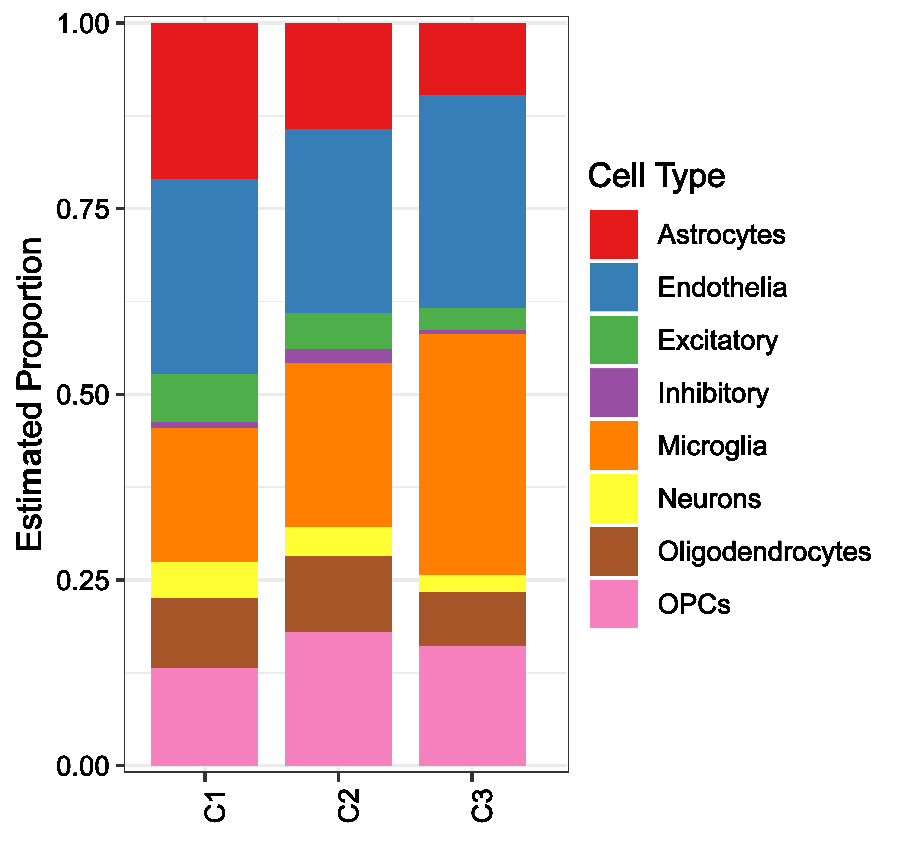

Fig. S5. RNA-seq analysis of BOs. Stacked graph of composition estimated in BOs (n = 3). Deconvolution was performed using CIBERSORT and the Cell Atlas signature.

Table S1. Primer sequences used for PCR analysis.

| **Species** | **Gene** | **Forward Primer（5'to3'）** | **Reverse Primer（5'to3'）** |
| --- | --- | --- | --- |
| Rat | *Actin* | AGATCAAGATCATTGCTCCTCCT | ACGCAGCTCAGTAACAGTCC |
|  | *Tjp1* | CGCAGCCACAAGCTATTCAT | CTTGTGATACGTGCGAGGTG |
|  | *Ocln* | CCCTTCTTTCCTTAGGCGACC | GTGCATCTCTCCGCCATACA |
|  | *Cldn5* | CTGCCTTAATGTCCGGTGGTAC | GTCCTTTGGTTCAGTAGGTTCCT |
|  | *Mfsd2a* | GCCACTCCCTGCTTCCATTAT | CAGGCACAAACCAGATGAGAAAG |
| Human | *ACTIN* | CCTTCCTGGGCATGGAGTC | TGATCTTCATTGTGCTGGGTG |
|  | *TJP1* | TCACGCAGTTACGAGCAAGT | TGAAGGTATCAGCGGAGGGA |
|  | *OCLN* | CCAATGTCGAGGAGTGGGTTAAA | AGTCATCCACAGGCGAAGTTAAT |
|  | *CLDN5* | CTGCTGCCTTACTTCCCAGAG | AAGCAGATTCTTAGCCTTCCCAC |
|  | *MFSD2A* | GCCTTGTTTCCAGGACCTCAATA | CTGGGCTTCATAGGGTTCTCTCT |
|  | *CAV-1* | GACCCTAAACACCTCAACGATGA | CCAGATGTGCAGGAAAGAGAGAA |
|  | *IL-10* | CAAGCCTTGTCTGAGATGATCCA | TGTCAAACTCACTCATGGCTTTG |
|  | *IL-1β* | CAGAAGTACCTGAGCTCGCC | AGATTCGTAGCTGGATGCCG |
|  | *IL-6* | ACCCCCAGGAGAAGATTCCA | GATGCCGTCGAGGATGTACC |
|  | TNFα | GCTGCACTTTGGAGTGATCG | TCACTCGGGGTTCGAGAAGA |
|  | *IBA1* | TGAGAAGACTGGTGGGAGAGAAG | GTTGATCTCATCCAGCCTCTCTT |
|  | *GFAP* | GCACGCAGTATGAGGCAATG | TAGTCGTTGGCTTCGTGCTT |
|  | *TMEM119* | TCCAGGGTCAGATTACAAGAGCAC | ACTGTTGATTCTGGAGGGTTTGA |
| *P. gingivalis* | 16S rRNA | GCGTGAAGGAAGACTGTCCTA | ACTCCCCAGGTGGATTACTTA |
|  | primer 1 | AGCTTGCCATACTGCGACTGAC | GATGTGGGTTGCGCTCGTTATG |
|  | primer 2 | GGCGTGGGTATCAAACAGGA | AAACCACATGTTCCTCCGCT |

Table S2. RT-qPCR reaction condition Ⅰ

| **Temperature** | **Time** | **Cycle Number** |
| --- | --- | --- |
| 95 °C | 10 min | 1 |
| 95 °C | 10 s | 40 |
| 60 °C | 30 s |  |
| 72 °C | 30 s |  |
| 90 °C | 10 s | 1 |
| 65 °C to 95 °C | 65℃ 5 s, 95℃ 0.5℃ | 1 |

Table S3. siRNA sequence.

| **Gene** | **sense（5'to3'）** | **antisense（5'to3'）** |
| --- | --- | --- |
| Cav-1 170 | CUGAGAAGCAAGUGUAUGATT | UCAUACACUUGCUUCUCAGTT |
| Cav-1 551 | GCAAGAUAUUCAGCAACAUTT | AUGUUGCUGAAUAUCUUGCTT |
| Cav-1 467 | GCUUCCUGAUUGAGAUUCATT | UGAAUCUCAAUCAGGAAGCTT |
| Cav-1 244 | GACGUGGUCAAGAUUGACUTT | AGUCAAUCUUGACCACGUCTT |
| NC | UUCUCCGAACGUGUCACGUTT | ACGUGACACGUUCGGAGAATT |

Table S4. Antibodies used in this study.

| **Antibodies** | **Dilution ratio** | **Cata number** | **RRID** |
| --- | --- | --- | --- |
| Anti-Cav-1 | 1: 3000 | Abcam, ab32577 | AB_725987 |
| Anti-β-actin | 1: 3000 | Abcam, ab8226 | AB_306371 |
| Anti-GFAP (WB) | 1: 10000 | Abcam, ab68428 | AB_1209224 |
| Anti-Iba1 (WB) | 1: 1000 | Abcam, ab178846 | AB_2636859 |
| Anti-MFSD2A | 1: 1000 | Abclonal, A22116 | AB_3739500 |
| HRP Anti-rabbit IgG （H+L） | 1: 10000 | Boster, BA1054 | AB_2734136 |
| HRP Anti-mouse IgG （H+L） | 1: 10000 | Boster, BA1050 | AB_2904507 |
| Anti-CD31 | 1: 100 | Abcam, ab76533 | AB_1523298 |
| Anti-CD13 | 1: 100 | Abcam, ab108310 | AB_10866195 |
| Anti-GFAP (IF) | 1: 150 | Abcam, ab68428 | AB_1209224 |
| Anti-ZO-1 | 1: 100 | Invitrogen, 402200 | AB_2533456 |
| Anti-Iba1 (IF) | 1: 100 | Abcam, ab178846 | AB_2636859 |
| Anti-rabbit secondary antibody (Alexa Fluor® 488) | 1: 200 | Abcam, ab150077 | AB_2630356 |
| Anti-mouse secondary antibody (Alexa Fluor® 488) | 1: 200 | Abcam, ab150113 | AB_2576208 |
| Anti-p-Tau | 1: 5000 | Abcam, ab254409 | AB_2905609 |
| Anti-Aβ | 1: 1000 | Abcam, ab201061 | AB_2722492 |

Table S5. Nest PCR amplification step one.

| **Temperature** | **Time** | **Cycle Number** |
| --- | --- | --- |
| 95 °C | 10 min | 1 |
| 95 °C | 15 s | 20 |
| 64 °C | 20 s |  |
| 72 °C | 9 s |  |
| 72 °C | 5 min | 1 |

Table S6. Nest PCR amplification step two.

| **Temperature** | **Time** | **Cycle Number** |
| --- | --- | --- |
| 95 °C | 10 min | 1 |
| 95 °C | 30 s | 40 |
| 60 °C | 30 s |  |
| 72 °C | 1 min |  |
| 72 °C | 5 min | 1 |

**Table S7. The chromatographic gradient of mobile phase (A: water, and B: acetonitrile).**

| **Min** | **Flow rate (mL/min)** | **A (%)** | **B (%)** |
| --- | --- | --- | --- |
| Initial | 0.6 | 90 | 10 |
| 0.3 | 0.6 | 90 | 10 |
| 2.0 | 0.6 | 10 | 90 |
| 2.5 | 0.6 | 10 | 90 |
| 2.6 | 0.6 | 90 | 10 |
| 3.5 | 0.6 | 90 | 10 |

**Table S8. NCBI alignment results for PCR band sequencing data.**

| **Sample** | | **Scientific Name** | **Max Score** | **Total Score** | **Query Cover** | **E value** |
| --- | --- | --- | --- | --- | --- | --- |
| *P. gingivalis* | 1 | *Porphyromonas gingivalis W83* | 852 | 3409 | 99% | 0.0 |
|  | 2 | *Porphyromonas gingivalis W83* | 841 | 3365 | 99% | 0.0 |
|  | 3 | *Porphyromonas gingivalis W83* | 843 | 3372 | 100% | 0.0 |
|  | 4 | *Porphyromonas gingivalis W83* | 865 | 3461 | 99% | 0.0 |
| 2VO + *P. gingivalis* | 1 | *Porphyromonas gingivalis W83* | 865 | 3461 | 99% | 0.0 |
|  | 2 | *Porphyromonas gingivalis W83* | 859 | 3439 | 99% | 0.0 |
|  | 3 | *Porphyromonas gingivalis W83* | 861 | 3446 | 99% | 0.0 |
